# In silico design of a multi-epitope vaccine against tick-borne encephalitis virus via immunoinformatic analysis

**DOI:** 10.1101/2024.10.26.620446

**Authors:** Bingjie Wang

**Affiliations:** State Key Laboratory of Virology College of Life Sciences Wuhan University Wuhan, Hubei, China

**Keywords:** ***K*eywords** Tick-borne encephalitis virus *·* Immunoinformatics *·* Multi-epitope vaccine *·* Molecular docking *·* Immune simulation

## Abstract

Tick-borne encephalitis virus (TBEV) is a serious pathogen that poses a significant threat to humans, causing encephalitis that can result in lifelong sequelae. In this study, we focused on the complete proteomes of the five current TBEV subtypes to identify dominant epitopes. Immunoinformatics tools were employed to screen for LBL, HTL, and CTL epitopes. These epitopes were then linked using various linkers and combined with adjuvants and histidine tag. The vaccine underwent a series of physicochemical property analyses, including secondary structure prediction, three-dimensional structure prediction, molecular docking, molecular dynamics simulation, immune simulation, and in silico cloning. The results indicate that the vaccine is highly conserved, strongly immunogenic, stable, non-allergenic, and non-toxic. Molecular docking and molecular dynamics simulation demonstrate that the vaccine can form a stable binding complex with TLR3. Immune simulation analysis shows that the vaccine effectively stimulates both cellular and humoral immune responses, accompanied by an increase in cytokine titers. Furthermore, through codon optimization and in silico cloning, the vaccine can be stably and effectively expressed in the *Escherichia coli* system. As an effective candidate for TBEV vaccination, the multi-epitope vaccine developed in this study has promising application prospects and provides a new approach for the research, development, and improvement of vaccines targeting TBEV.

## 1 Introduction

Tick-borne encephalitis virus (TBEV) is a single-stranded, positive-sense, enveloped RNA virus (Pulkkinen et al., 2018). According to reports from the International Committee on Taxonomy of Viruses (ICTV), it belongs to the Flaviviridae family and the *Orthoflavivirus* genus. Other viruses within this genus include Powassan virus (POWV), dengue virus (DENV), Zika virus (ZIKV), Japanese encephalitis virus (JEV), West Nile virus (WNV), and yellow fever virus (YFV) (Simmonds et al., 2017). In nature, TBEV infects both ticks and various wild animals, with ticks serving as the primary transmission vector. The virus is transmitted to humans primarily through tick bites, although transmission can also occur via breast milk, laboratory exposure, and blood transfusions (Lindquist and Vapalahti, 2008; Süss, 2011; Avšič-Županc et al., 1995; Ličková et al., 2021). Humans serve as the terminal host for TBEV infection. TBEV poses a significant threat to human health, contributing to over 10,000 reported cases globally each year (Chiffi et al., 2023). Furthermore, the incidence of TBEV infections has notably increased in Europe in recent years (Kwasnik et al., 2023).

The genome size of tick-borne encephalitis virus (TBEV) is approximately 11,000 base pairs (bp) and contains a single open reading frame, which encodes a precursor protein. This precursor is subsequently cleaved by both viral and host cell proteases into three structural proteins (C, prM, and E) and seven non-structural proteins (NS1, NS2A, NS2B, NS3, NS4A, NS4B, and NS5) (Füzik et al., 2018; Pulkkinen et al., 2018). The viral envelope is embedded with prM and E proteins, with prM cleavage being essential for TBEV maturation (Pierson and Diamond, 2020). The E protein, which forms dimers, entirely covers the virus particle’s surface and is involved in receptor binding, fusion, and cellular invasion. E serves as the primary target of host humoral immunity (Agudelo et al., 2021). The geographical distribution of TBEV is closely linked to that of its tick vectors and is primarily found in Eurasia (Im et al., 2020). In recent years, global climate change has contributed to an expansion in the distribution range of ticks, resulting in the emergence of TBEV in new regions and populations, thereby increasing potential threats (De Graaf et al., 2016). TBEV is classified into three main subtypes: European (TBEV-Eu), Siberian (TBEV-Sib), and Far Eastern (TBEVFE). Among these, TBEV-FE exhibits the highest virulence, followed by TBEV-Sib, while TBEV-Eu has the lowest virulence (Tonteri et al., 2013). Additionally, ongoing research has identified two other subtypes of TBEV: Baikal (TBEV-Bkl) and Himalayan (TBEV-Him) (Dai et al., 2018; Sukhorukov et al., 2023).

Currently, although there are approved vaccines available, all are in the form of inactivated virus vaccines and exhibit a single formulation (Kollaritsch et al., 2012). Furthermore, these vaccines were introduced in the last century and have not been updated for decades. Consequently, there is an urgent need to develop new TBEV vaccines to address the increasingly severe threat posed by TBEV. Our research aims to employ immunoinformatics tools to design a multi-epitope vaccine against TBEV that can effectively activate an immune response. To encompass the five TBEV subtypes, we selected epitopes that are highly conserved, non-immunogenic, non-toxic, and highly antigenic across these subtypes. These sequences are linked using specialized linkers to create a multi-epitope vaccine. We then evaluated the antigenicity, allergenicity, and physicochemical properties of this vaccine. Following this, we predicted its three-dimensional structure and performed molecular docking with TLR3. The results of molecular dynamics simulation indicate that the multi-epitope vaccine protein can bind naturally to TLR3. Finally, we conducted immune simulation, which demonstrated that the multi-epitope vaccine effectively stimulates an immune response. In conclusion, these findings suggest that the multi-epitope vaccine we designed possesses superior characteristics and may serve as an effective candidate for controlling TBEV-related health risks in the population.

## 2 Results

### 2.1 Selection of linear B-cell epitopes (LBL)

Epitopes with a length of 16 mer and a score greater than 0.8 were selected. Subsequently, epitopes present in all five subtypes were further identified and considered conserved. We then evaluated the antigenicity, allergenicity, and toxicity of these epitopes, retaining only those that were highly antigenic, non-allergenic, and non-toxic. This process yielded one epitope from the C protein, four from the E protein, five from the NS3 protein, four from the NS5 protein, while no suitable epitopes were identified in other proteins (Table S1). Finally, based on the positions and score magnitudes of these epitopes in the TBEV proteome, at least one epitope was retained for each position, resulting in a total of six epitopes (Table 1). Given that the E protein is located on the surface of the virus particle and serves as a natural target for neutralizing antibodies, three epitopes from the E protein were retained.

**Table 1:**
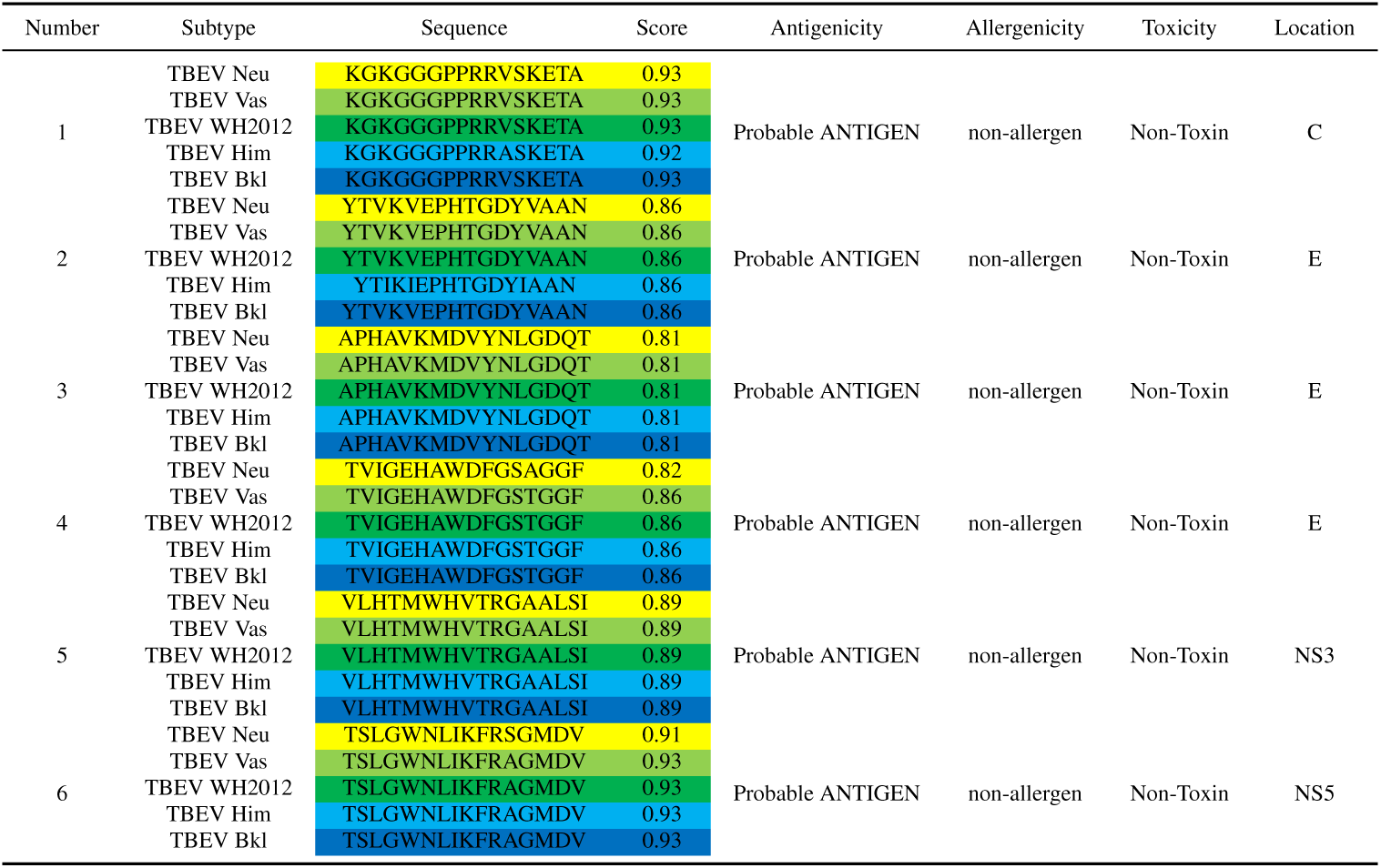
Summary of selected LBL epitopes.

### 2.2 T-cell epitope prediction

T-cell epitopes must bind to MHC complexes to participate in the activation of cellular immunity. Numerous HLA alleles express MHC complexes, and the binding patterns of these complexes vary across different alleles, leading to polypeptide specificity. Considering that TBEV is primarily distributed in Eurasia, we first identified the HLA alleles with high frequencies in the Eurasian population, followed by the prediction of T-cell epitopes based on these alleles. Based on the research conducted by Sanchez-Mazas et al. (2024), we selected HLA alleles with high distribution frequencies in the Eurasian population, ensuring that at least one allele from each subtype was included (Table S2).

Cytotoxic T-lymphocyte epitopes (CTL) with a length of 9 mer were predicted. Only those epitopes with high affinity, present in all five subtypes, and exhibiting high antigenicity, non-allergenicity, and non-toxicity were retained, resulting in a total of eight epitopes (Table S3). These epitopes were distributed across the prM, NS2A, NS3, NS4A, and NS5 proteins. Based on the HLA genotypes and affinities corresponding to these epitopes, at least one epitope was retained for each genotype, yielding a total of six epitopes (Table 2).

**Table 2:**
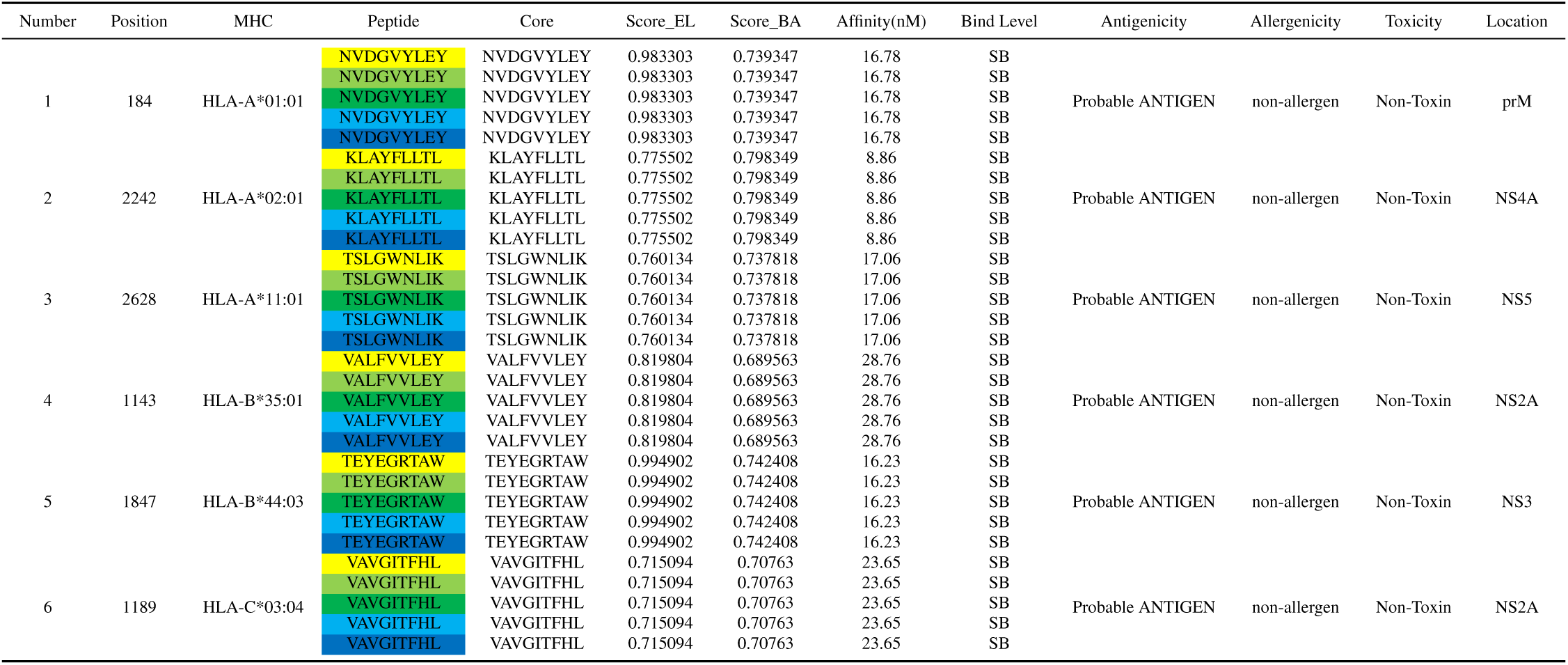
Summary of selected CTL epitopes.

Helper T-lymphocyte epitopes (HTL) with a length of 15 mer were predicted. Only those epitopes with high affinity, present in all five subtypes, and exhibiting high antigenicity, non-allergenicity, strong induction of interferon-*γ*, and non-toxicity were retained, resulting in a total of nine epitopes (Table S4). These epitopes were distributed across the NS1, NS3, NS4B, and NS5 proteins. Based on the HLA genotypes and affinities corresponding to these epitopes, at least one epitope was retained for each genotype, yielding a total of six epitopes (Table 3).

**Table 3:**
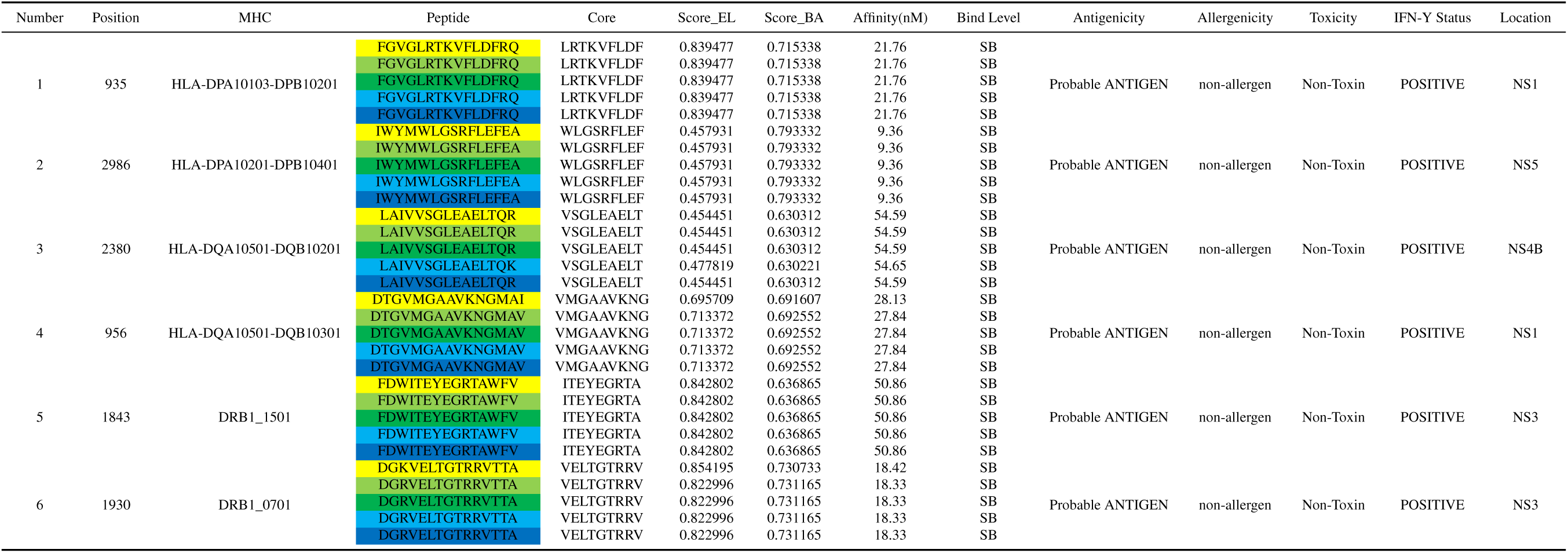
Summary of selected HTL epitopes.

### 2.3 Population coverage of selected epitopes

The examination results of HTL and CTL epitopes indicated that the global coverage of CTL epitopes was 72.33%, while the global coverage of HTL epitopes was 97.53% (Figure 2, S1). Given that the multi-epitope vaccine contains both types of epitopes, integrating them provides an accurate measure of overall population coverage. The results showed that the multi-epitope vaccine achieved a population coverage of 99.32%. Since we previously selected HLA alleles with high frequencies in the Eurasian population for T-cell epitope prediction, it is theoretically expected that the population coverage of the multi-epitope vaccine in Europe would be relatively high, which was confirmed by the results. Specifically, the coverage of CTL epitopes in Europe was 79.24%, while the coverage of HTL epitopes was 99.94%, both exceeding the global population coverage. Notably, the coverage of CTL epitopes in the UK reached 85.54% (Table S5).

**Figure 1:**
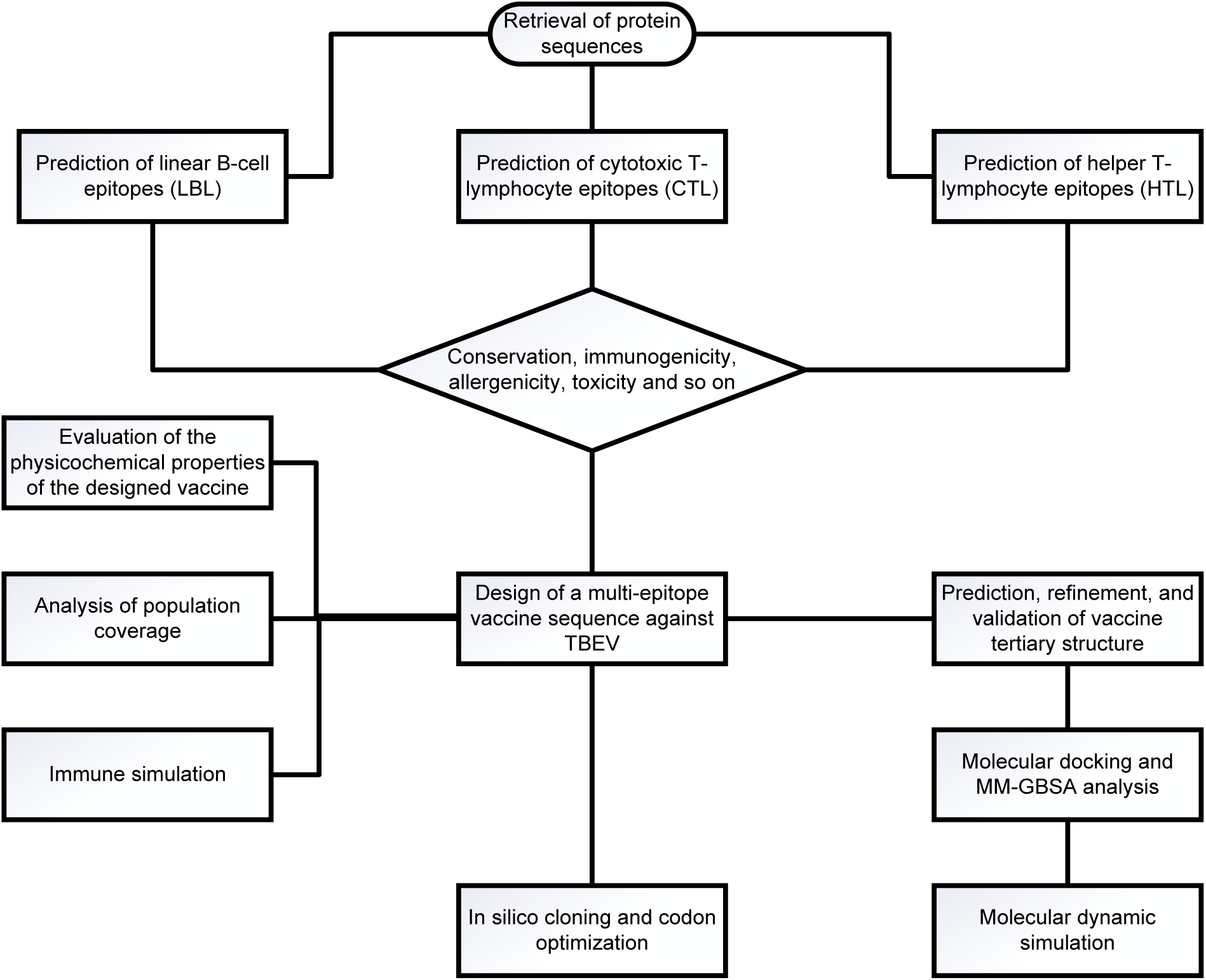
Research flowchart of the multi-epitope vaccine against TBEV.

**Figure 2:**
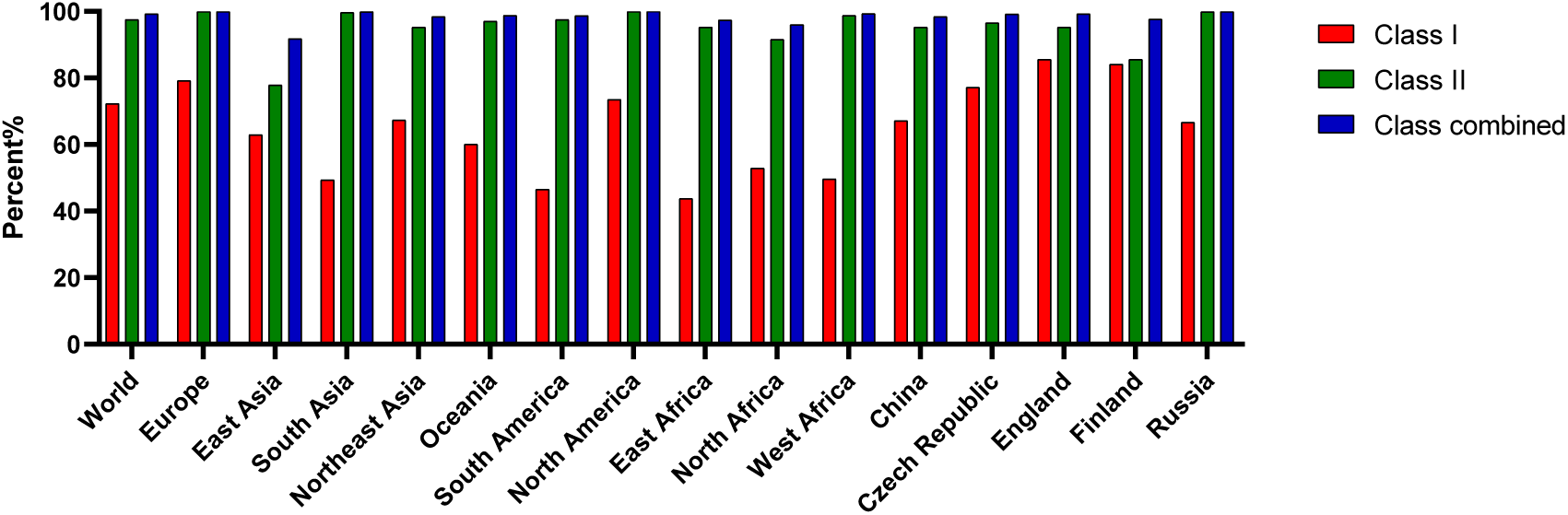
The population coverage of the selected epitopes across different countries and ethnicities.

**Figure 3:**
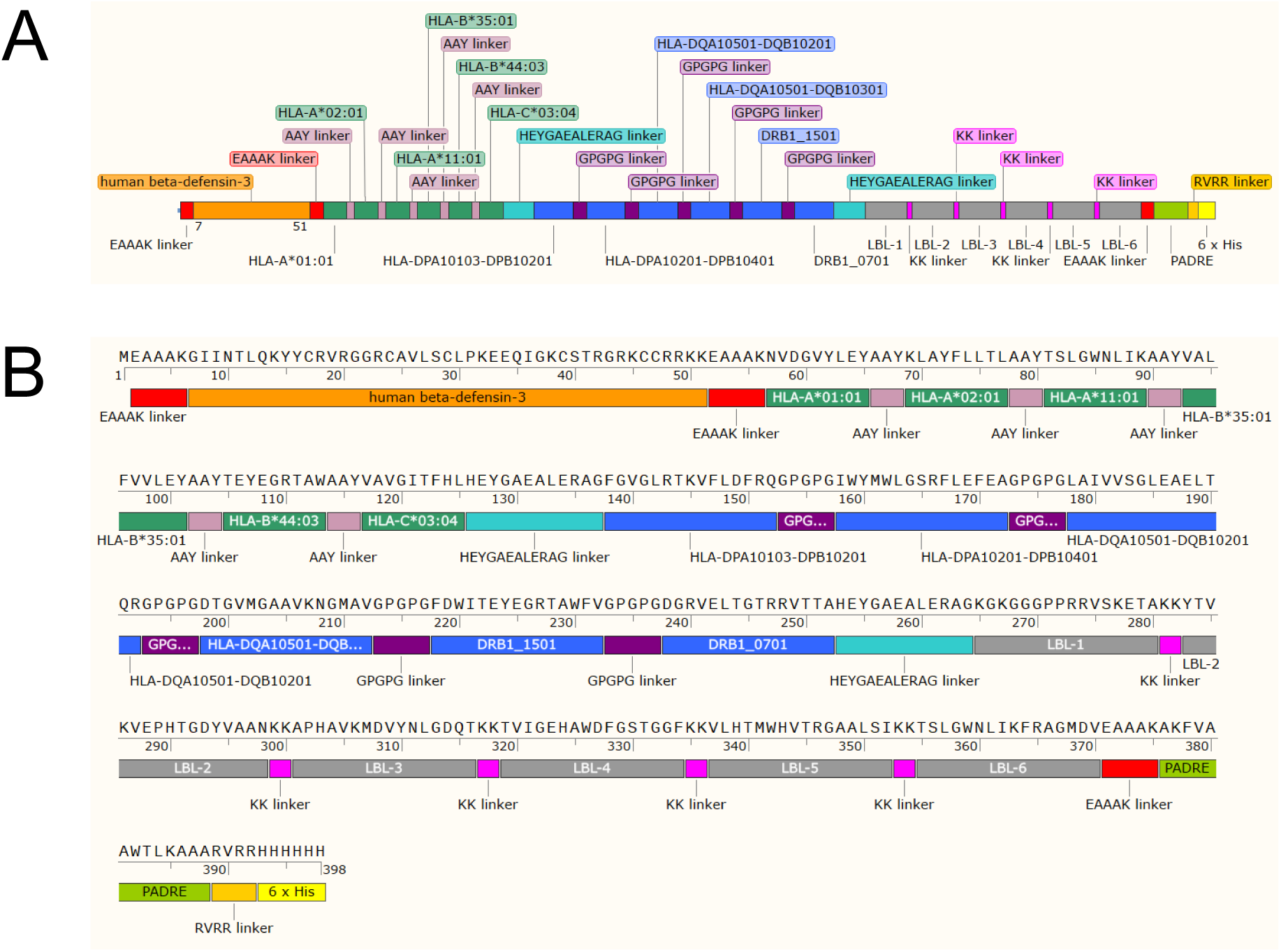
Topological representation of the constructed vaccine. (**A**) The overall topology of the vaccine. (**B**) Protein sequence information of the vaccine.

### 2.4 Construction of the multi-epitope vaccine

The final selected epitopes were linked using specific linkers. The AAY linker was chosen to connect MHC-I epitopes, while the GPGPG linker was employed for MHC-II epitopes. The human beta-defensin-3 and PADRE sequences, serving as adjuvants, were linked at the N-terminus through the EAAAK linker. The last MHC-I epitope was connected to the first MHC-II epitope via the HEYGAEALERAG linker. Additionally, the KK linker was utilized to combine B-cell epitopes. A histidine tag was incorporated at the C-terminus and linked to the final construct through the RVRR linker (Figure 2).

### 2.5 Evaluation of the multi-epitope vaccine

The multi-epitope vaccine comprises 398 amino acids and has a molecular weight of 43.3 kDa (Table 4). Its instability index is 18.19, indicating that it is stable. The GRAVY value is -0.235, classifying it as a hydrophilic protein. The vaccine has a half-life of over 10 hours in *Escherichia coli* and is soluble, which facilitates protein expression and purification. The vaccine protein does not possess a transmembrane helix or a signal peptide (Figure 4A, 4B). Its antigenicity score is 0.6620, and it exhibits no allergenicity or toxicity. The secondary structure of the vaccine protein is illustrated in the figure (Figure 4C). Overall, the multi-epitope vaccine we designed demonstrates excellent physicochemical properties.

**Table 4:** Physical and chemical features of the multi-epitope vaccine.

| Number | Physicochemical Properties | Results |
| --- | --- | --- |
| 1 | Total amino acids residue | 398 |
| 2 | Molecular weight | 43339.86 |
| 3 | Instability index | 18.19 (Stable) |
| 4 | GRAVY value | -0.235 |
| 5 | Estimated half-life (mammalian reticulocytes, in vitro), (yeast, in vivo), (Escherichia coli, in vivo) | 30 hours, >20 hours, >10 hours |
| 6 | Theoretical pI | 9.63 |
| 7 | Antigenicity | 0.6620 ( Probable ANTIGEN ) |
| 8 | Allergenicity | non-allergen |
| 9 | Toxicity | non-toxin |
| 10 | Solubility and usability of proteins expressed in E. coli | 0.4714 (solubility) |

**Figure 4:**
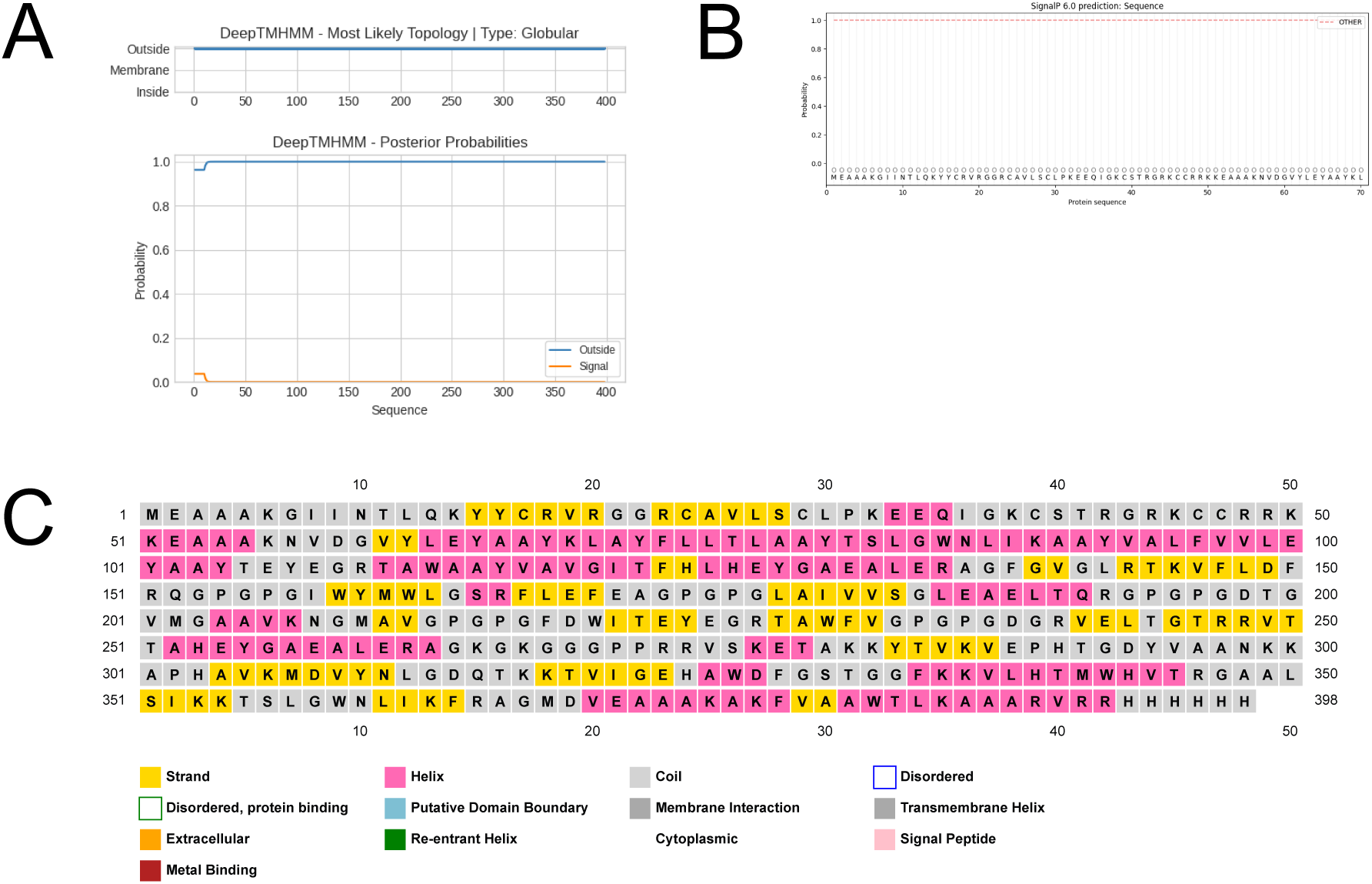
Results of candidate vaccine evaluation. (**A**) Prediction of transmembrane domains. (**B**) Prediction of signal peptide. (**C**) Prediction of secondary structure.

### 2.6 Structure modeling and validation of the multi-epitope vaccine

We utilized the Robetta server for protein structure prediction, generating five models. Subsequently, we refined these five models using the Galaxy server, resulting in a total of 25 models. We then conducted a structural assessment of these models. Based on the evaluations from three different tools, model 4.1 was determined to be the best (Figure 5A, Table S6). The Z-score of this model was -6.81, and the quality factor calculated by ERRAT was 91.545 (Figure 5C). Analysis of the Ramachandran plot indicated that 88.8% of the amino acids in the model were in favored regions, with only 0.9% in disallowed regions (Figure 5B). We proceeded with model 4.1 for subsequent experiments.

**Figure 5:**
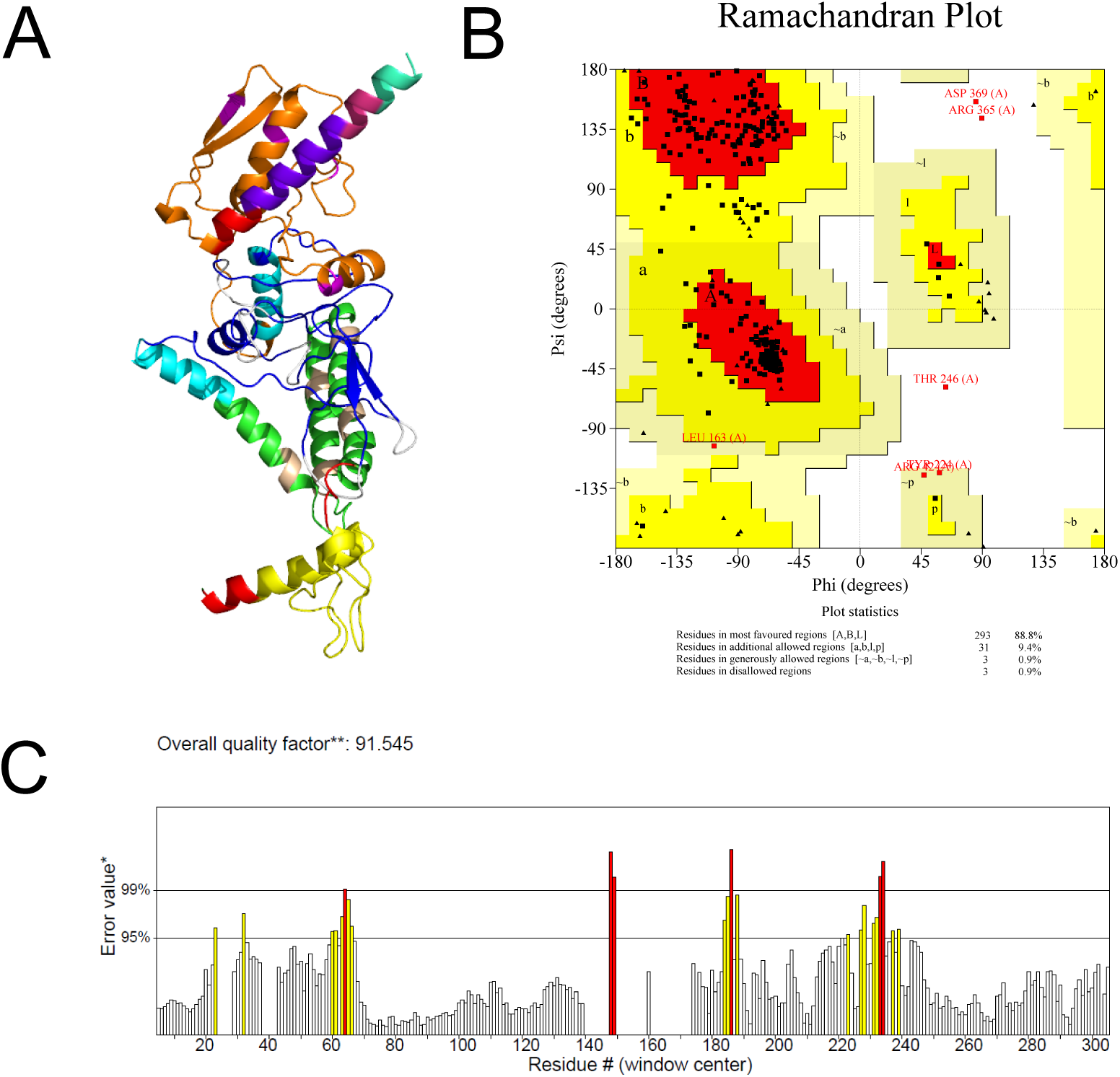
Modeling and evaluation of the three-dimensional structure of the multi-epitope vaccine. (**A**) The three-dimensional structure of the vaccine. Different components of the vaccine are represented by different colors. (**B**) Ramachandran plot for all amino acids of the multi-epitope vaccine. (**C**) Displays the ERRAT score, which is used to evaluate the quality of the refined model. Regions with error rates exceeding 95% are highlighted in yellow, while residues with error rates surpassing 99% are extremely rare and are marked in red. CD4^+^ T cells (Th1 and Th2) and CD8^+^ T cells (CTL) (Figure 8C, 8D, S3, S4). The candidate vaccine also induced immune cells to produce IFN-*γ* and a series of cytokines, resulting in three peaks (Figure 8F). These findings suggest that after three rounds of immune stimulation, the simulation demonstrated a strong induction of an effective immune response. Overall, the simulated immune response indicates that this vaccine can induce both cellular and humoral immunity, making it a promising candidate.

### 2.7 Docking of candidate vaccine with TLR3 and molecular dynamics analysis

We employed two prediction tools for docking to mutually validate our findings. Among the top ten models predicted by HDOCK, one model was highly similar to one of the top ten models predicted by pyDockWEB (Figure 6A, 6B). Consequently, we believe that these two docking modes may closely represent the actual interaction. A detailed analysis of the interaction interface revealed that the interface in the vaccine protein was almost identical in both docking modes (Figure 6C, 6D). In TLR3, the differences between the two docking modes were primarily located at both ends of the binding region. Each end contained a helix: the helix at the lower end participated in the binding for the model predicted by HDOCK, while the upper end did not. Conversely, in the model predicted by pyDockWEB, the helix at the upper end participated in the binding, while the lower end did not. We then conducted molecular dynamics simulations on these two models. The RMSD results indicated that within 50 ns, the HDOCK docking model had values ranging from 0.25 to 1.5, while the pyDockWEB docking model ranged from 0.25 to 1.2 (Figure 7A, 7C). This suggests that the docking mode of the pyDockWEB model is more stable. Additionally, the results of the MM-GBSA analysis showed that the binding free energy of the HDOCK model was -34.46 kcal/mol, while that of the pyDockWEB model was -54.4 kcal/mol. The lower binding free energy of the pyDockWEB model further confirms the molecular dynamics simulation results, indicating that its docking mode is superior.

**Figure 6:**
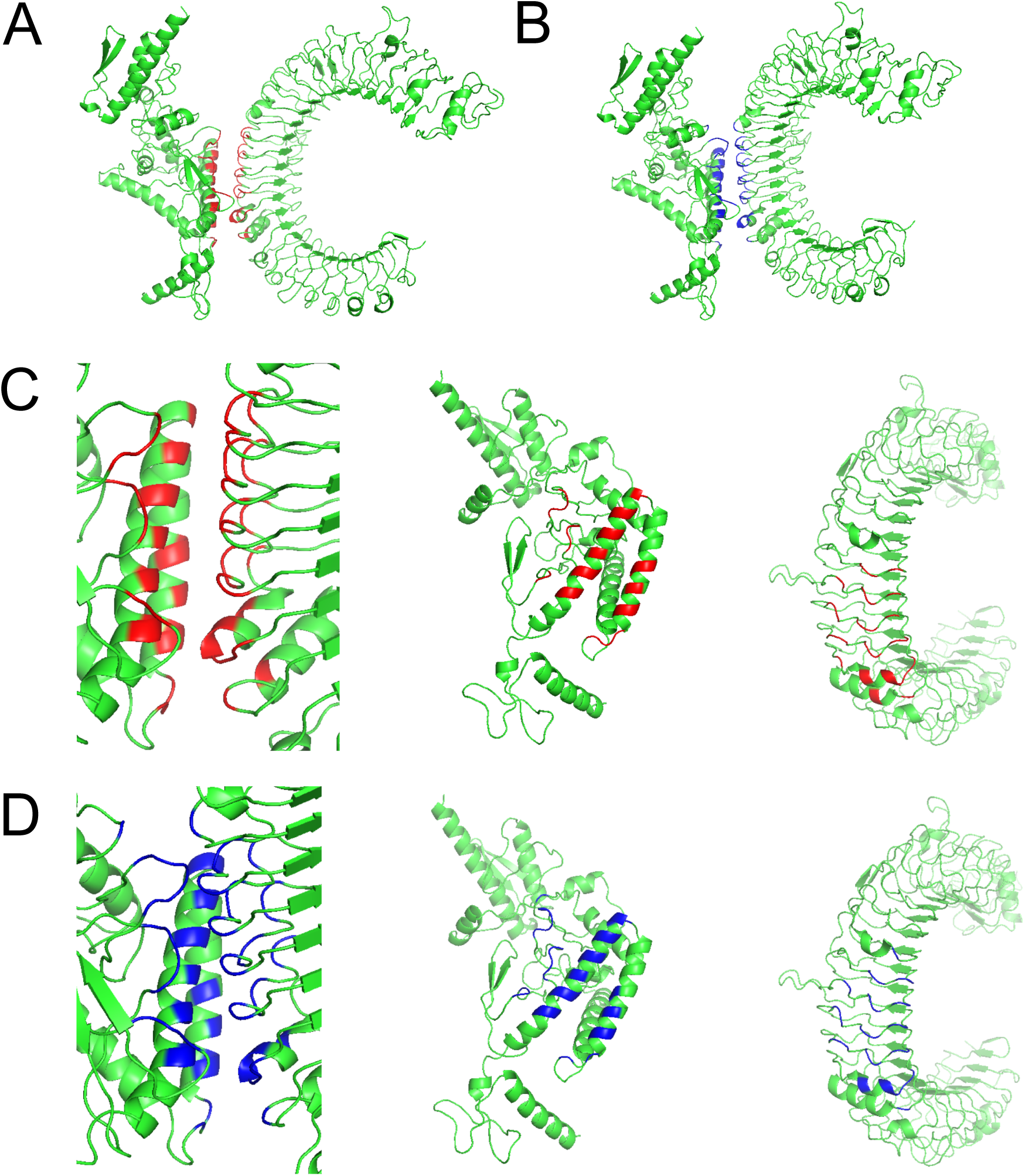
Docking results of candidate vaccine with TLR3 molecule. (**A**) The docking complex predicted by HDOCK. (**B**) The docking complex predicted by pyDockWEB. (**C**) The binding pattern of the docking complex predicted by HDOCK. (**D**) The binding pattern of the docking complex predicted by pyDockWEB.

**Figure 7:**
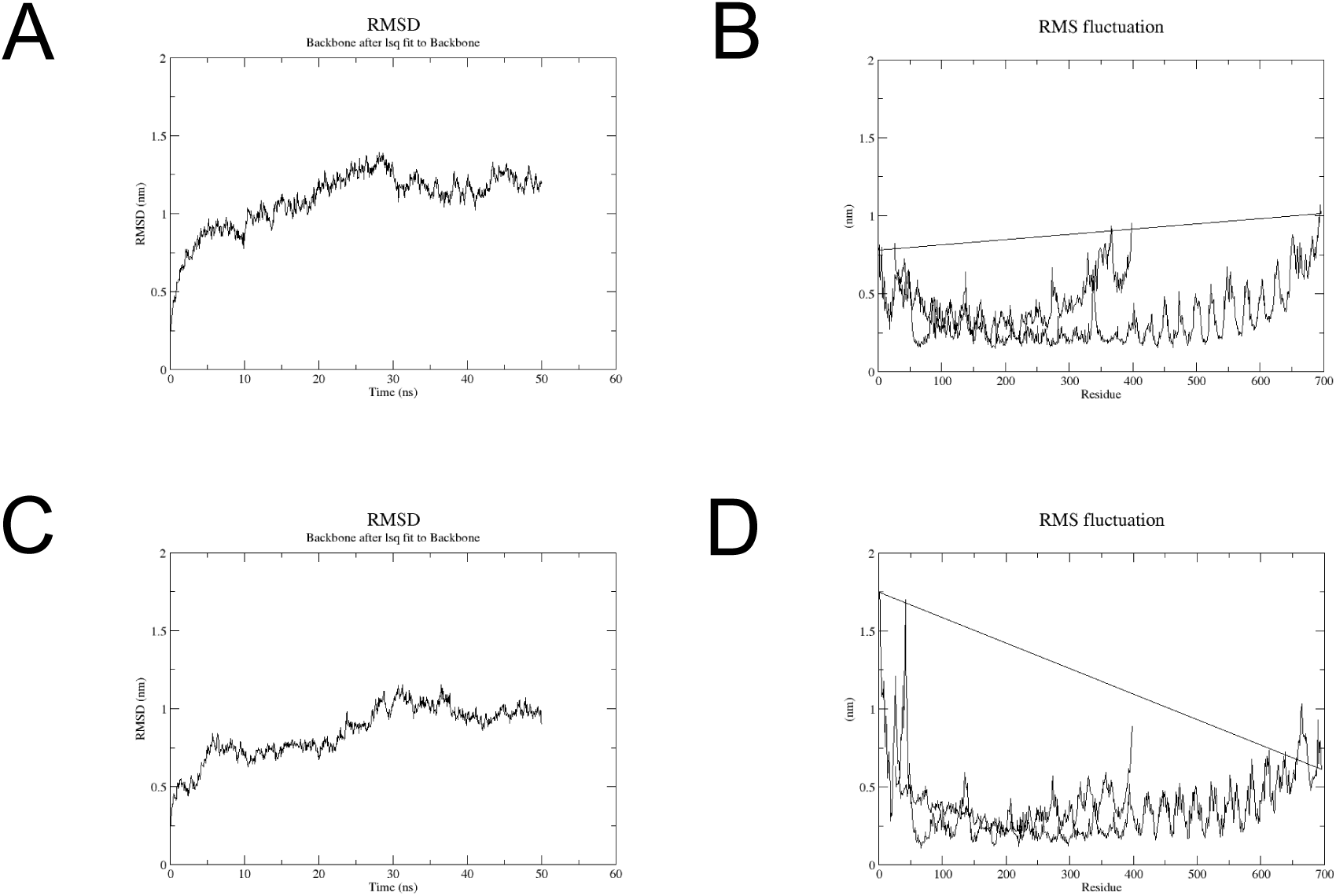
The results of molecular dynamics simulation of the vaccine-TLR3 complex. (**A**) The RMSD analysis of the docking complex predicted by HDOCK. (**B**) The RMSD analysis of the docking complex predicted by pyDock-WEB. (**C**) The RMSF analysis of the docking complex predicted by HDOCK. (**D**) The RMSF analysis of the docking complex predicted by pyDockWEB.

### 2.8 Immune simulation of the vaccine

The intensities of both specific and non-specific immune responses are crucial in the vaccine immunization process. We conducted an immune simulation of the candidate vaccine, and the results indicated that the vaccine protein could activate NK, DC, and macrophage (MA) cells (Figure 8E, S5). Additionally, antigen-presenting cells (APCs) stimulated the differentiation and proliferation of B lymphocytes, leading to a significant increase in B cell populations and levels of IgG1, IgG2, IgM, and IgG + IgM antibodies (Figure 8A, 8B, S2). An upward trend was also observed in

**Figure 8:**
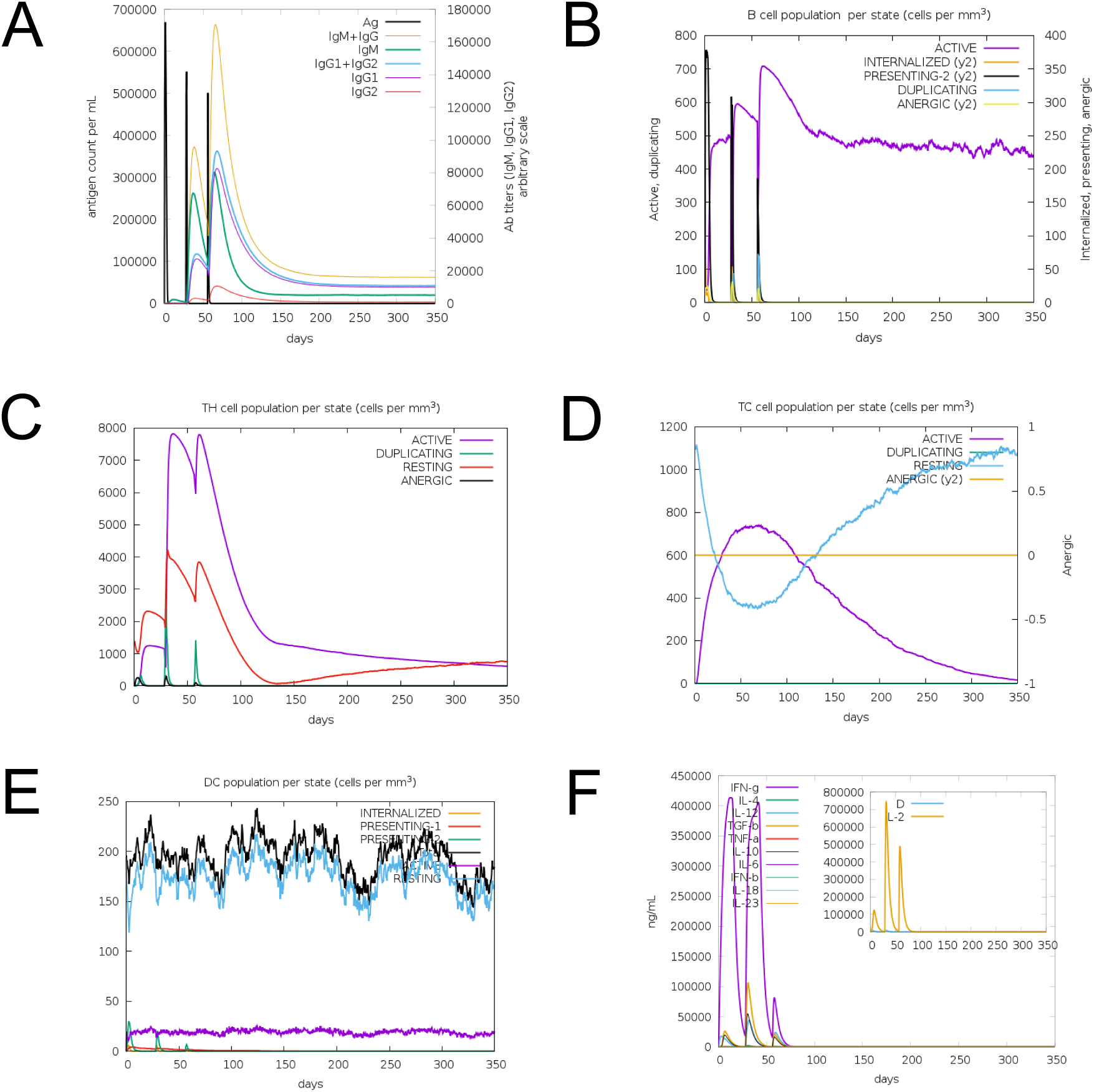
Immune simulation of the vaccine designed against TBEV. (**A**) The change in antibody titer after vaccine immunity. (**B**) The change in the number of B cells after vaccine immunity. (**C**) The change in the number of TH cells after vaccine immunity. (**D**) The change in the number of TC cells after vaccine immunity. (**E**) The change in the number of DC cells after vaccine immunity. (**F**) The changes of the secondary immune elements response after vaccine immunity.

### 2.9 In silico cloning of the multi-epitope vaccine in the pET-21a expression vector

To maximize the expression of the multi-epitope vaccine in the Escherichia coli expression system, we performed codon optimization. The codon adaptation index (CAI) value of the optimized vaccine sequence was 0.79, and the GC content was 56.11%. The CAI value falls within the appropriate range for high expression, while the ideal GC content is between 30% and 70%. These results indicate that the optimized vaccine is likely to be expressed efficiently in Escherichia coli. The full-length optimized sequence is presented in Table S7. Upon completion of the optimization, we successfully cloned the full-length vaccine sequence into the pET-21a expression vector using the restriction sites Nde I and Xho I (Figure 9).

**Figure 9:**
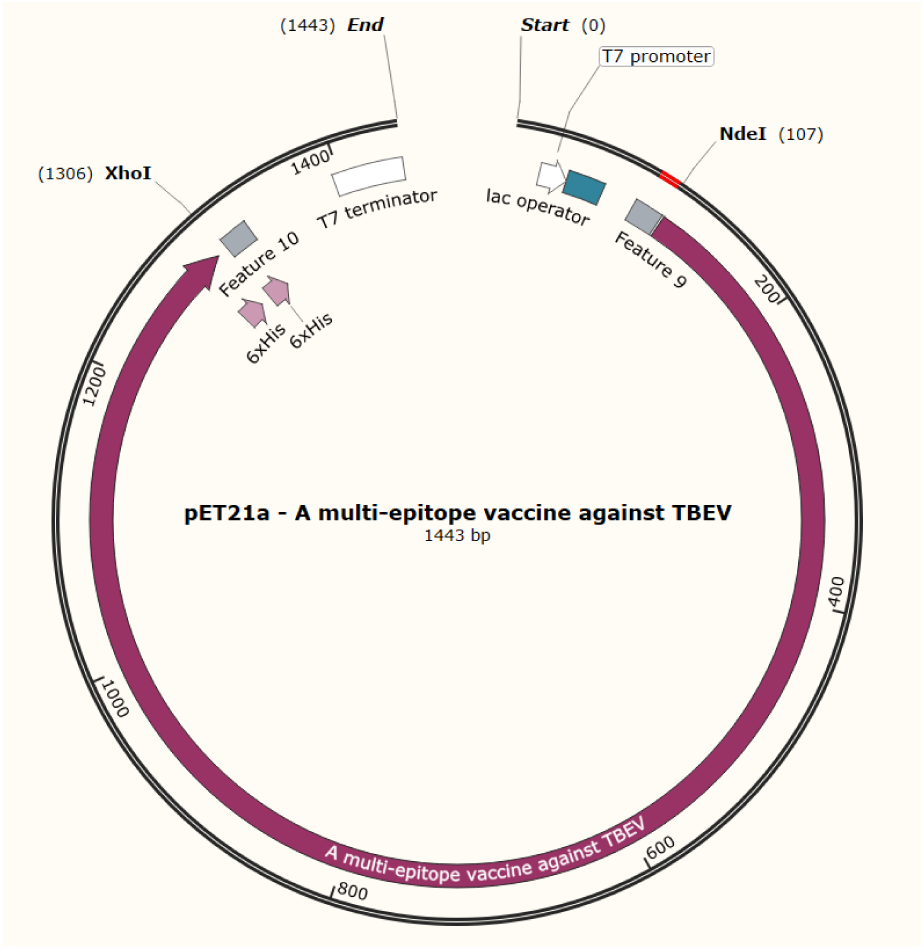
Construction of the multi-epitope vaccine expression system. Candidate vaccine inserted into pET-21a.

## 3 Discussion

Despite the availability of approved vaccines, TBEV infections continue to exhibit a significant upward trend in Europe, with instances of breakthrough infections reported (Andersson et al., 2010). The urgent development of new vaccines against TBEV is essential. Epitope-based vaccines can safely and effectively elicit immune responses by targeting conserved epitopes from various antigens. The application of immunoinformatics methods in the design of epitope-based vaccines has become increasingly prevalent, as these approaches can significantly reduce resource and time expenditures in the development of vaccines against viruses and bacteria (Alharbi et al., 2024; Rani et al., 2024; Tarrahimofrad et al., 2021).

In the proteome analysis of TBEV, the sequences AAA86870.1, AAD34205.1, AIG51207.1, ABS00285.1, and AWK48861.1 represent the five current subtypes of the virus. These polyproteins were utilized for subsequent epitope prediction and vaccine design. It is well established that viral mutations can diminish vaccine efficacy; therefore, designing a vaccine that is effective against all TBEV subtypes is crucial for developing a multi-epitope vaccine. Compared to single-peptide-based vaccines, epitope-based vaccines often elicit a stronger immune response. To enhance the immunogenicity of the vaccine, we designed a multi-epitope-based vaccine that aims to elicit an immune response while minimizing the risk of allergic reactions. This epitope-based vaccine is primarily composed of highly conserved, non-allergenic, strongly antigenic, and non-toxic epitopes. The overall population coverage of these epitopes is projected to reach 99% globally, indicating that the vaccine can accommodate diverse ethnic groups. Predictions regarding the physicochemical properties of the vaccine suggest that it can be stably expressed in the Escherichia coli system, is hydrophilic, lacks a transmembrane helix and signal peptide, and can therefore be easily prepared and stably transported.

We conducted molecular docking studies of the vaccine and its receptor to analyze their binding interface, as this interaction is crucial for eliciting an immune response against a specific pathogen. According to the docking results, the designed vaccine can effectively bind to the targeted TLR3 receptor. Furthermore, the stability of the docking complex is essential for generating a long-lasting and sustained immune response against the pathogen. Molecular dynamics simulation is a computer-based method used to analyze the physical movements of the docking complexes and evaluate their dynamic behavior. The results of the molecular dynamics simulation indicate that stable intermolecular interactions and binding conformations suggest the vaccine has significant potential to stimulate a robust immune response. The RMSD analysis predicts that the vaccine and the receptor molecule maintain appropriate binding in a dynamic environment, enabling effective activation of the immune system. Additionally, through MM-GBSA analysis, we observed a binding free energy of -54.4 kcal/mol, further confirming the vaccine’s ability to bind effectively to TLR3. In conclusion, both molecular docking and molecular dynamics simulations demonstrate that the vaccine exhibits stable binding with TLR3, allowing the immune system to readily recognize it and generate an appropriate immune response.

Cytokines are signaling molecules essential for a robust immune response. High titers of cytokines were observed in the multi-epitope vaccine we constructed. Additionally, the results of the immune simulation demonstrate that the vaccine can activate both humoral and cellular immunity. Therefore, in the context of vaccine development against TBEV, our vaccine emerges as an excellent candidate.

## 4 Conclusion

In summary, we have successfully designed a multi-epitope vaccine against TBEV by applying advanced bioinformatics, immunoinformatics, and various biochemical methods, including molecular docking and molecular dynamics simulation. For epitope prediction, we utilized the proteomes of five TBEV subtypes as input. Following this prediction, we further screened the identified epitopes through immunoinformatics analysis and selected six potential epitopes corresponding to LBL, HTL, and CTL. Using these epitopes, we designed a multi-epitope vaccine. The immunoinformatics analysis of the vaccine indicated its significant potential to activate a robust immune response against TBEV. Molecular docking and molecular dynamics simulation predicted that the vaccine can stably bind to TLR3, suggesting that it can effectively induce an immune response in the host. Given our prediction that this vaccine can elicit an immune response against TBEV, in vivo experimental studies are necessary to further confirm its immunogenic efficacy.

## 5 Materials and methods

The methodology of this study is illustrated in Figure 1.

### Retrieval of protein sequences

Currently, there are five subtypes of tick-borne encephalitis virus (TBEV). The proteome sequences of these subtypes can be retrieved from the National Center for Biotechnology Information (NCBI) (Sayers et al., 2022). They are as follows: European (TBEV Neu, GenBank: AAA86870.1), Siberian (TBEV Vas, GenBank: AAD34205.1), Far Eastern (TBEV WH2012, GenBank: AIG51207.1), Baikal (TBEV Bkl, GenBank: ABS00285.1), and Himalayan (TBEV Him, GenBank: AWK48861.1).

### Prediction of linear B-cell epitopes (LBL)

B cells play a crucial role in the immune system by generating a long-lasting humoral immune response. Consequently, B-cell epitopes are among the most important factors in vaccine design. In this study, we utilized the ABCpred server (https://webs.iiitd.edu.in/raghava/abcpred/ABC_submission.html) (Saha and Raghava, 2006) to identify linear B-cell epitopes present in all five subtypes of tick-borne encephalitis virus (TBEV). For ABCpred, we adjusted the length parameter to 16 and set the threshold to 0.8. ABCpred is a B-cell epitope prediction service based on artificial neural networks and covers a wide range of pathogen groups, including viruses, bacteria, fungi, and protozoa. Based on the scoring, we selected epitopes with high scores. All selected epitopes will undergo further evaluation for immunogenicity, toxicity, and allergenicity using the VaxiJen 2.0 server (https://ddg-pharmfac.net/vaxijen/VaxiJen/VaxiJen.html) (Doytchinova and Flower, 2007), the ToxinPred 3.0 server (https://webs.iiitd.edu.in/raghava/toxinpred3/) (Gupta et al., 2013), and the AllerTOP v. 2.0 server (https://www.ddg-pharmfac.net/AllerTOP/) (Dimitrov et al., 2013), respectively. The AllerTOP v. 2.0 server employs automatic cross-covariance (ACC) and the k-nearest neighbor algorithm (k = 1) to predict protein sensitization based on factors such as hydrophobicity, molecular weight, secondary structure properties, and relative amino acid abundance. The final selection comprises those epitopes with an immunogenicity score greater than 0.25 that are non-toxic and non-allergenic.

### Prediction of cytotoxic T-lymphocyte epitopes (CTL)

The NetMHCpan-4.1 server (https://services.healthtech.dtu.dk/services/NetMHCpan-4.1/) (Reynisson et al., 2020) employs artificial neural networks (ANN) to predict the binding of peptides to any MHC molecule with a known sequence. This method is trained on a dataset comprising over 850,000 quantitative binding affinities (BA) and mass spectrometry-eluted ligand (EL) peptides. Epitopes are evaluated based on their IC50 values, where a low IC50 value indicates strong binding to MHC class I molecules. The screening criterion selects a threshold of IC50 < 100 nM, as this represents high-affinity binding between the epitope and the MHC molecule, suggesting significant potential for immune recognition. The predicted epitopes are subsequently evaluated for conservation, immunogenicity, allergenicity, and toxicity.

### Prediction of helper T-lymphocyte epitopes (HTL)

The NetMHCIIpan-4.3 server (https://services.healthtech.dtu.dk/services/NetMHCIIpan-4.3/) employs artificial neural networks (ANN) to predict the binding of peptides to HLA class II molecules. It is trained on an extensive dataset comprising over 650,000 binding affinity (BA) and eluted ligand mass spectrometry (EL) measurements, covering three human MHC class II isotypes: HLA-DR, HLA-DQ, and HLA-DP, as well as mouse (H-2) and bovine (BoLA-DRB3) molecules. Fifteen-mer epitopes are selected based on IC50 values. Predicted HTL epitopes with IC50 values below 100 nM are assessed for their capacity to induce interferon gamma using the IFNepitope server (https://webs.iiitd.edu.in/raghava/ ifnepitope/predict.php) (Dhanda et al., 2013). Epitopes that strongly induce interferon gamma require further evaluation for conservation, immunogenicity, allergenicity, and toxicity.

### Analysis of population coverage for predicted epitopes

HLA alleles exhibit varying expressions and distributions across different populations and regions, which can influence the effectiveness of epitope-based vaccines. Assessing population coverage is crucial for accurately predicting epitopes for different HLA bindings. Epitopes with enhanced HLA binding can accommodate varying frequencies across ethnic groups, thereby mitigating the MHC restriction of T cell responses. We utilized the population coverage tool on the IEDB server (http://tools.iedb.org/tools/ population/iedb_input) (Vita et al., 2019) to determine the population coverage of selected epitopes. The selection of HLA alleles was based on the HLA genotypes corresponding to the previously predicted epitopes.

### Design of a multi-epitope vaccine sequence against TBEV

To design the multi-epitope vaccine, the most effective T cell and B cell epitopes were concatenated using various linkers, adjuvants, and PADRE sequences. In this study, the adjuvant sequence employed was human beta-defensin-3. The PADRE sequence was incorporated into the vaccine design to enhance the immune response of cytotoxic T lymphocyte (CTL) epitopes (Sarkar et al., 2021). The EAAAK linker connected human beta-defensin-3 to the PADRE sequence, while the AAY linker bound the CTL epitopes to one another (Arai et al., 2001). The HEYGAEALERAG linker served as the interface between the CTL and helper T lymphocyte (HTL) epitopes. Additionally, the GPGPG linker connected the HTL epitopes, and the KK linker was used to join the B cell epitopes. Finally, the RVRR linker combined the vaccine with the HisTag sequence. These design strategies aimed to enhance the immune response against TBEV.

### Evaluation of the physicochemical properties of the designed vaccine

The ProtParam server (https://web.expasy.org/protparam/) (Hebditch et al., 2017) is utilized to evaluate various physicochemical properties of the vaccine, including the aliphatic index, isoelectric point, molecular weight, GRAVY value, instability index, and estimated half-life. Additionally, the immunogenicity, allergenicity, and toxicity of the vaccine must also be assessed. Transmembrane helices are predicted using the DeepTMHMM server (https://services.healthtech.dtu.dk/ services/DeepTMHMM-1.0/) (Ikeda et al., 2002), while potential signal peptide sequences are predicted on the SignalP-6.0 server (https://services.healthtech.dtu.dk/services/SignalP-6.0/) (Nielsen, 2017). The solubility of the vaccine expressed in Escherichia coli is predicted using the NetSolP-1.0 server (https://services.healthtech.dtu.dk/services/NetSolP-1.0/) (Thumuluri et al., 2022). For an ideal vaccine candidate, high antigenicity, absence of allergenicity, an extracellular topological structure, and other criteria must be met.

### Prediction of the secondary structure of the vaccine

The PSIPRED server (http://bioinf.cs.ucl.ac.uk/ psipred/) (Karypis, 2006) is utilized to predict the secondary structure of the vaccine protein. The PSIPRED Workbench provides a range of protein structure prediction methods.

### Prediction, refinement, and validation of vaccine tertiary structure

The Robetta server (https://robetta.bakerlab.org/) (Kim et al., 2004) utilizes comparative modeling to predict protein structures and is employed for three-dimensional structure modeling. The five predicted models are further refined using the GalaxyRefine server (https://galaxy.seoklab.org/cgi-bin/submit.cgi?type=REFINE) (Heo et al., 2013), generating an additional five models for each original prediction. The quality of all models will be evaluated and compared using the SAVES v6.1 server (https://saves.mbi.ucla.edu/) (Laskowski et al., 2006) and the ProSA-web server (https://prosa.services.came.sbg.ac.at/prosa.php/) (Wiederstein and Sippl, 2007) to identify the best prediction model.

### Molecular docking and MM-GBSA analysis

The binding interaction of the vaccine with human toll-like receptor-3 (TLR-3, PDB ID: 2A0Z) was analyzed using the HADDOCK2.4 server (https://rascar.science.uu.nl/ haddock2.4/) (Van Zundert et al., 2016) and the pyDockWEB server (https://life.bsc.es/pid/pydockweb/) (Jiménez-García et al., 2013). These two tools respectively docked ten binding modes with the highest scores. By comparing the prediction results from both tools, a common binding mode was identified, which served as the final result. To evaluate the binding free energy of the vaccine-TLR3 complex, we employed the MM/GBSA method, accessible via the HawDock server (http://cadd.zju.edu.cn/hawkdock/) (Weng et al., 2019).

### Molecular dynamic simulation

We conducted a molecular dynamics simulation of the vaccine-TLR3 complex to evaluate the stability of their binding using the WebGRO server (https://simlab.uams.edu/ProteinInWater/ index.html) (Paz et al., 2004). Water molecules were used to solvate the entire complex. The simulation was performed at a temperature of 300 K and a pressure of 1.0 bar for 50 ns, generating 1,000 frames for each simulation. All other parameters were set to their default values. The simulation parameters calculated included root mean square deviation (RMSD) and root mean square fluctuation (RMSF).

### In silico cloning and codon optimization

To enhance protein expression levels, codon optimization is widely employed. In this study, we utilized the ExpOptimizer server (https://www.novopro.cn/tools/ codon-optimization.html#:∼:text=ExpOptimiz) to convert the vaccine’s protein sequence into a DNA sequence suitable for the Escherichia coli (E. coli) expression system. Using SnapGene software, we cloned the vaccine DNA sequence into the pET-21a expression vector.

### Immune simulation

The immune simulation of the vaccine was conducted using the C-IMMSIM server (https://kraken.iac.rm.cnr.it/C-IMMSIM/index.php?page=1) (Rapin et al., 2010), which provides a prediction of immune interactions that closely resemble real-life scenarios. Considering the four-week interval between the first and second doses of the vaccine, the simulation step parameter was set to 1,050. For three injections, the time steps were established at 1, 84, and 168, with each time step representing 8 hours in real life. All other parameters were set to their default values in this study.

## Supplementary Figures

**Figure S1:**
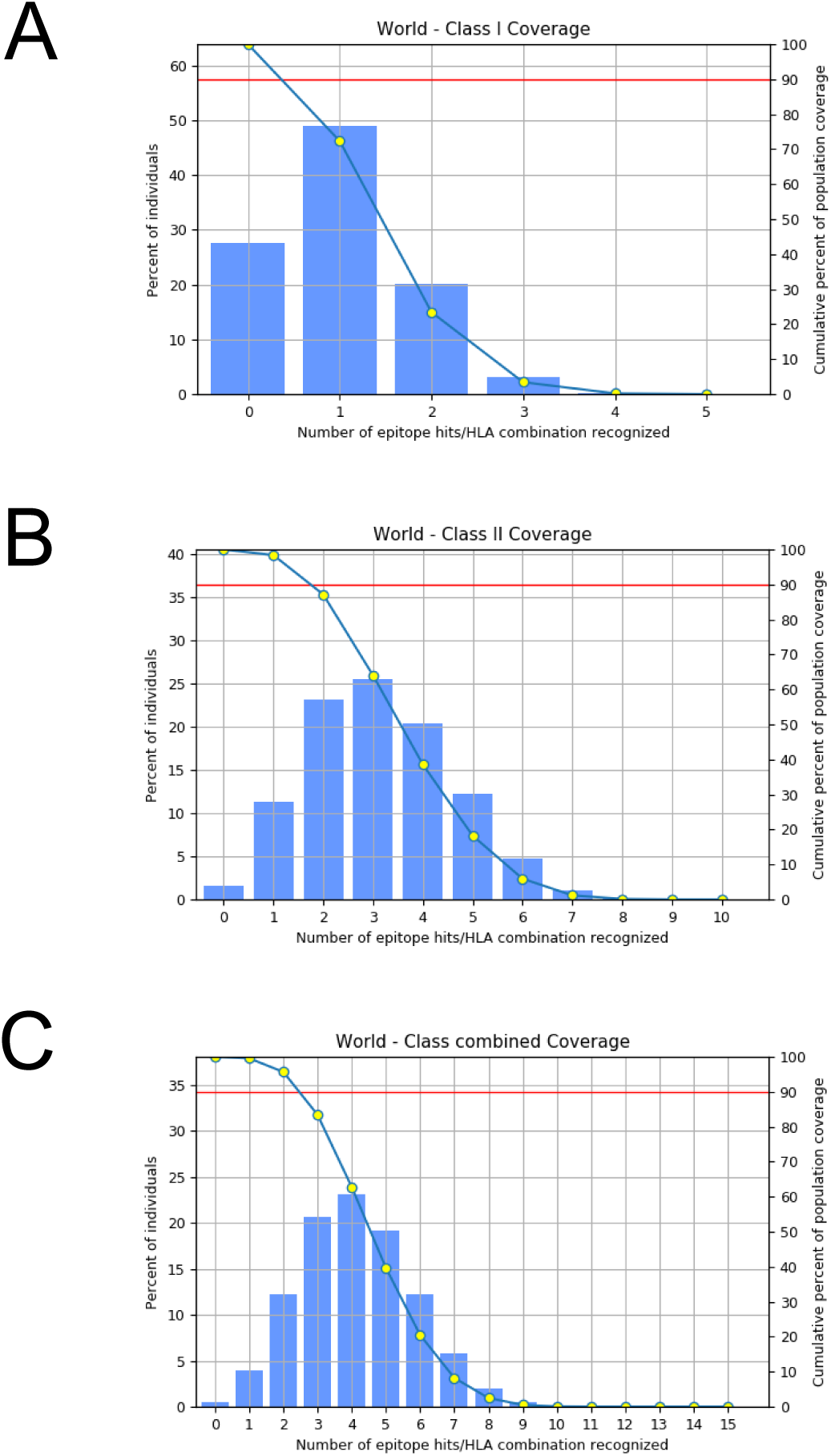
Global population coverage of CTL epitopes and HTL epitopes for both MHC-I and MHC-II. (**A**) MHC-I coverage. (**B**) MHC-II coverage. (**C**) Combined coverage.

**Figure S2:**
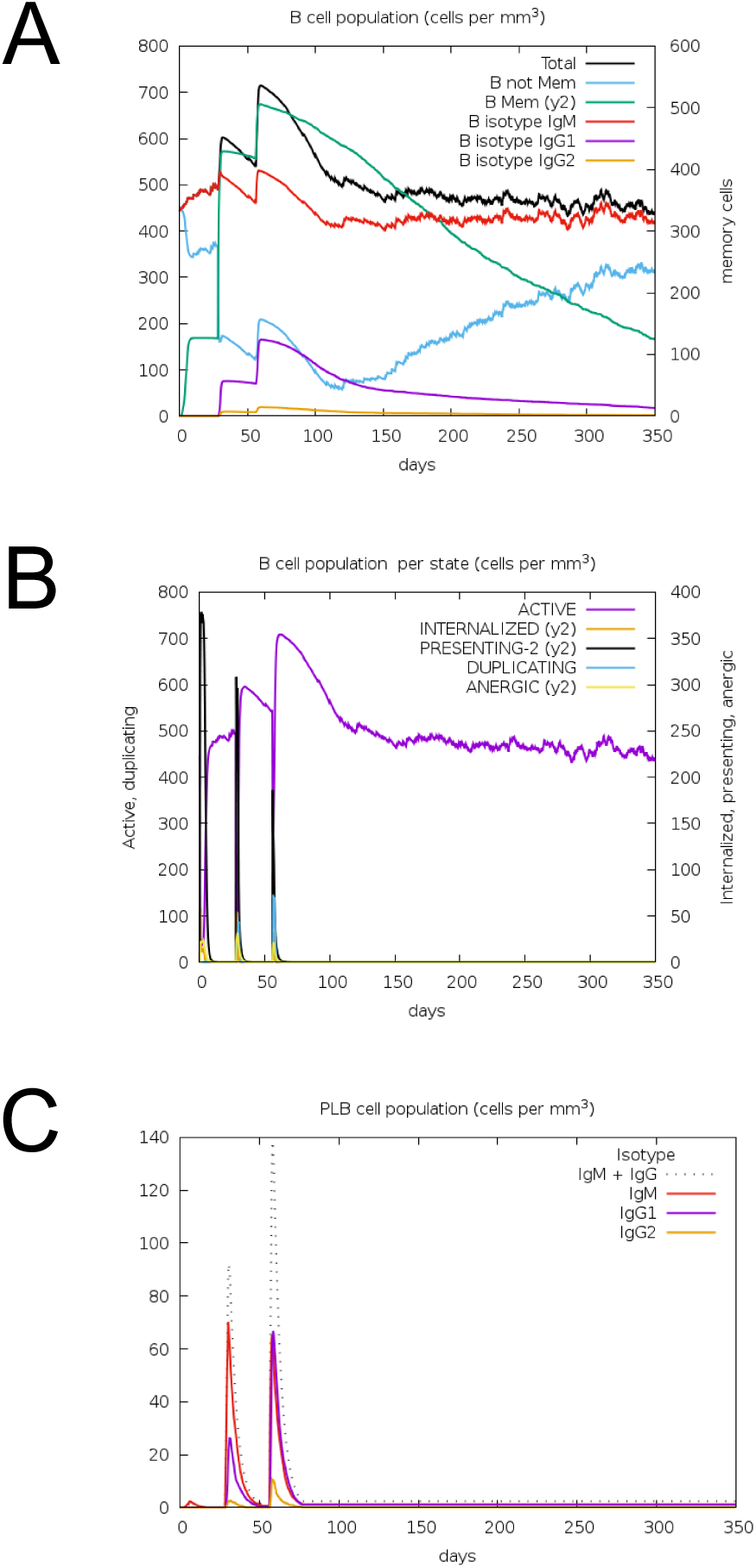
Display of changes in the number of B cells after vaccine immunity.

**Figure S3:**
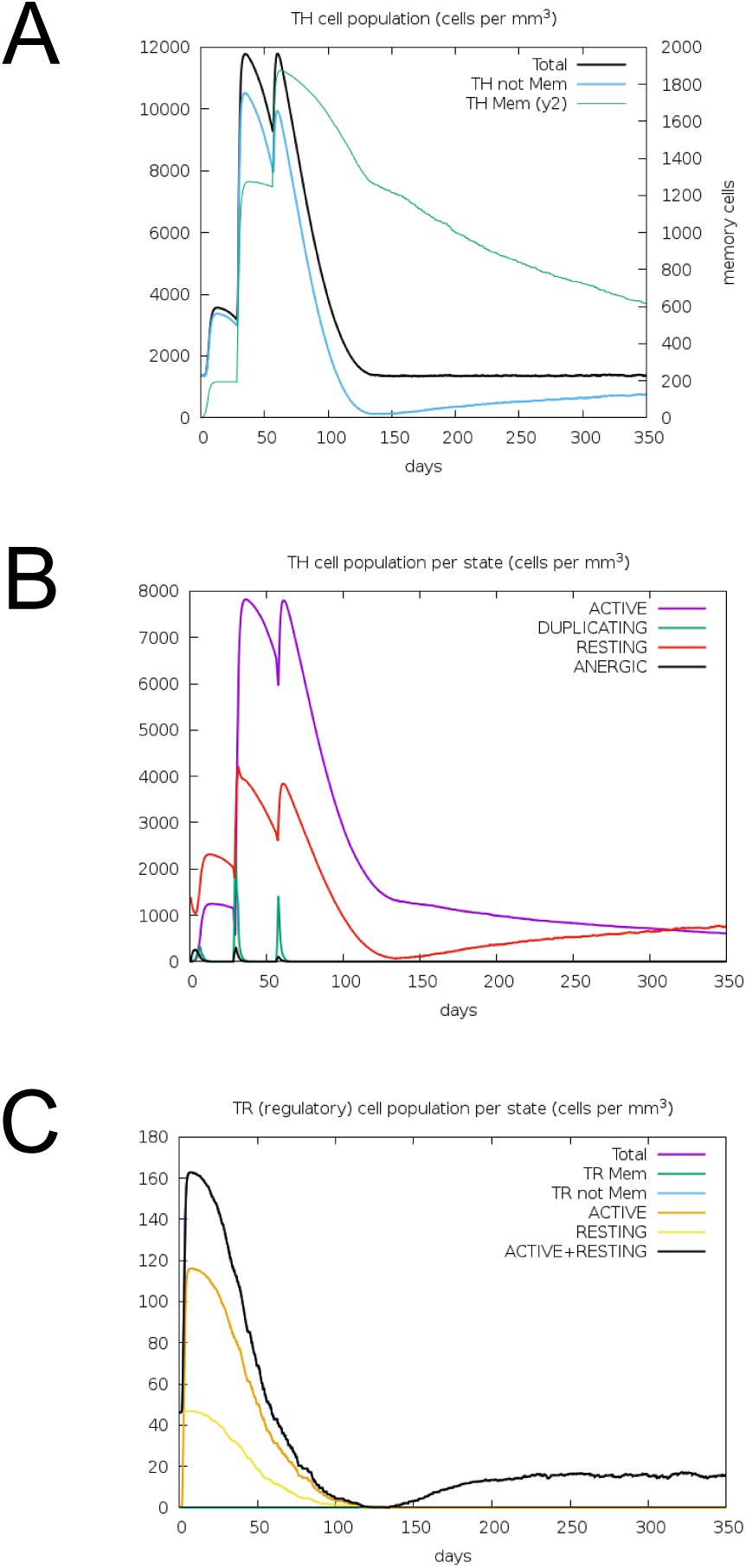
Display of changes in the number of TH cells after vaccine immunity.

**Figure S4:**
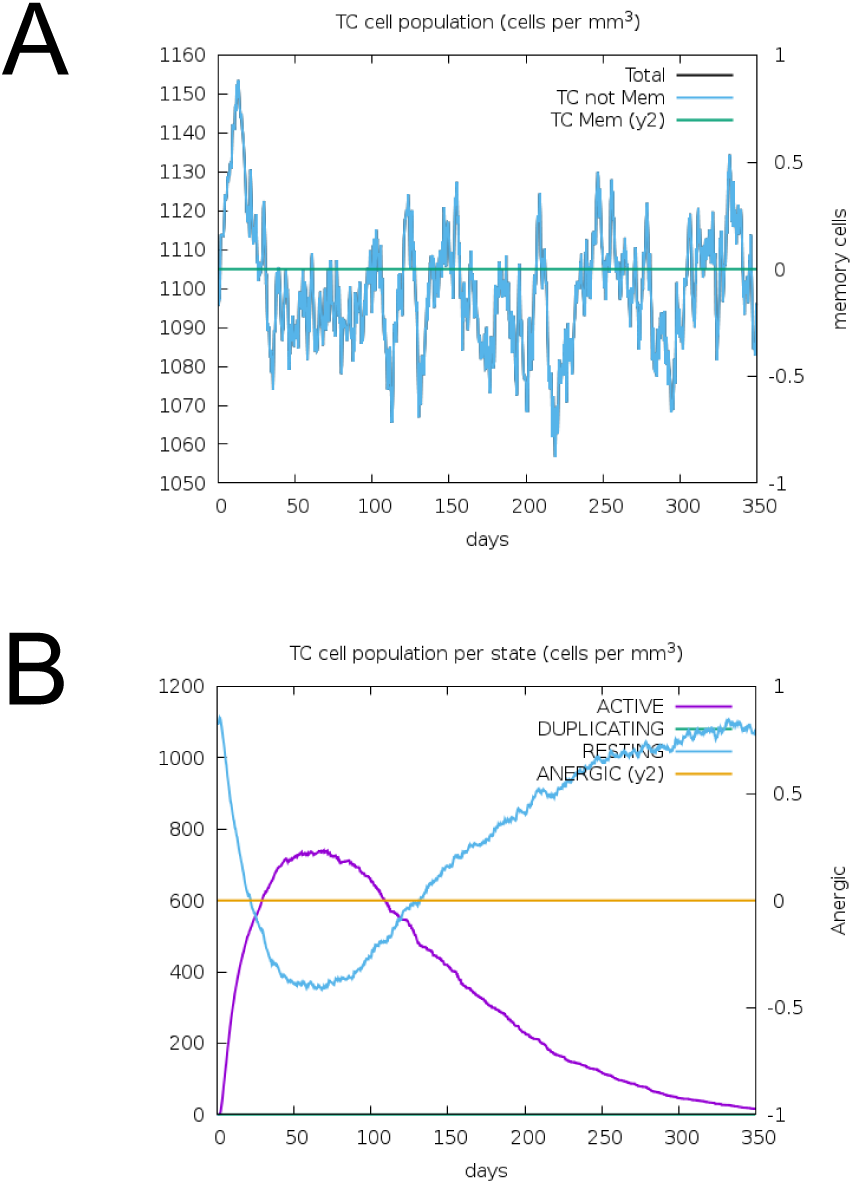
Display of changes in the number of TC cells after vaccine immunity.

**Figure S5:**
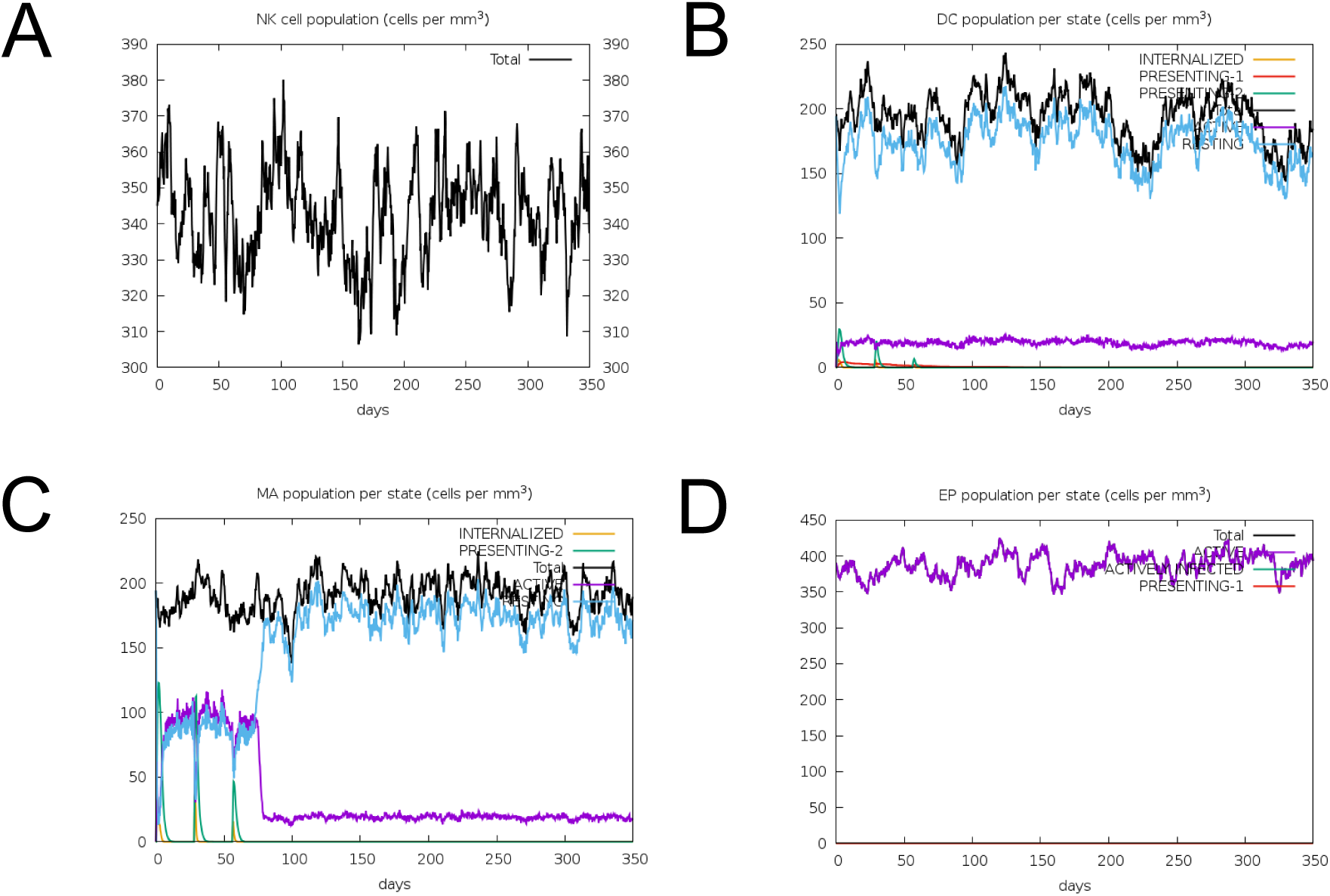
Display of changes in the number of NK, DC, MA and EP cells after vaccine immunity.

## Supplementary Tables

**Table S1:**
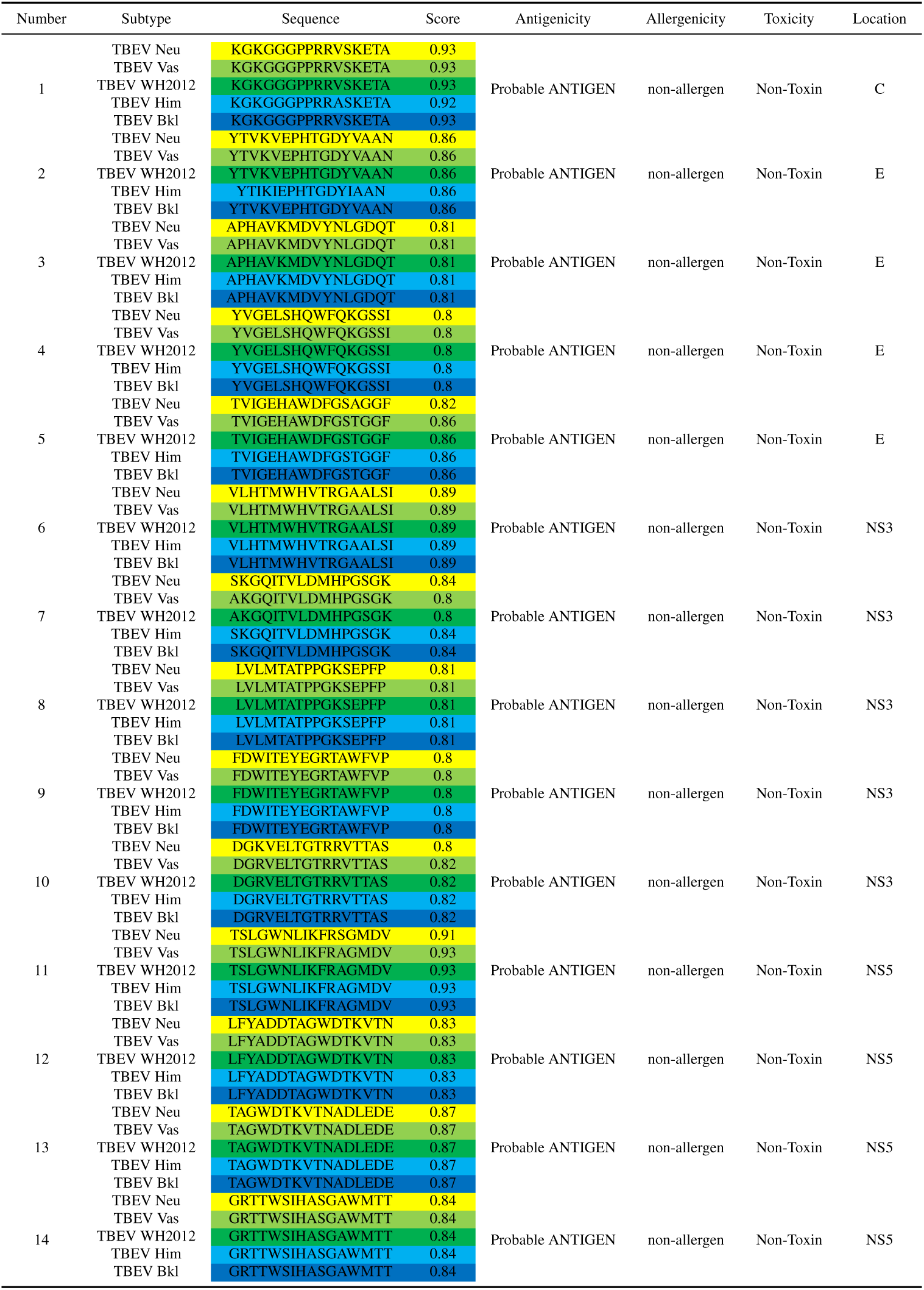
Summary of the screened LBL epitopes.

**Table S2:**
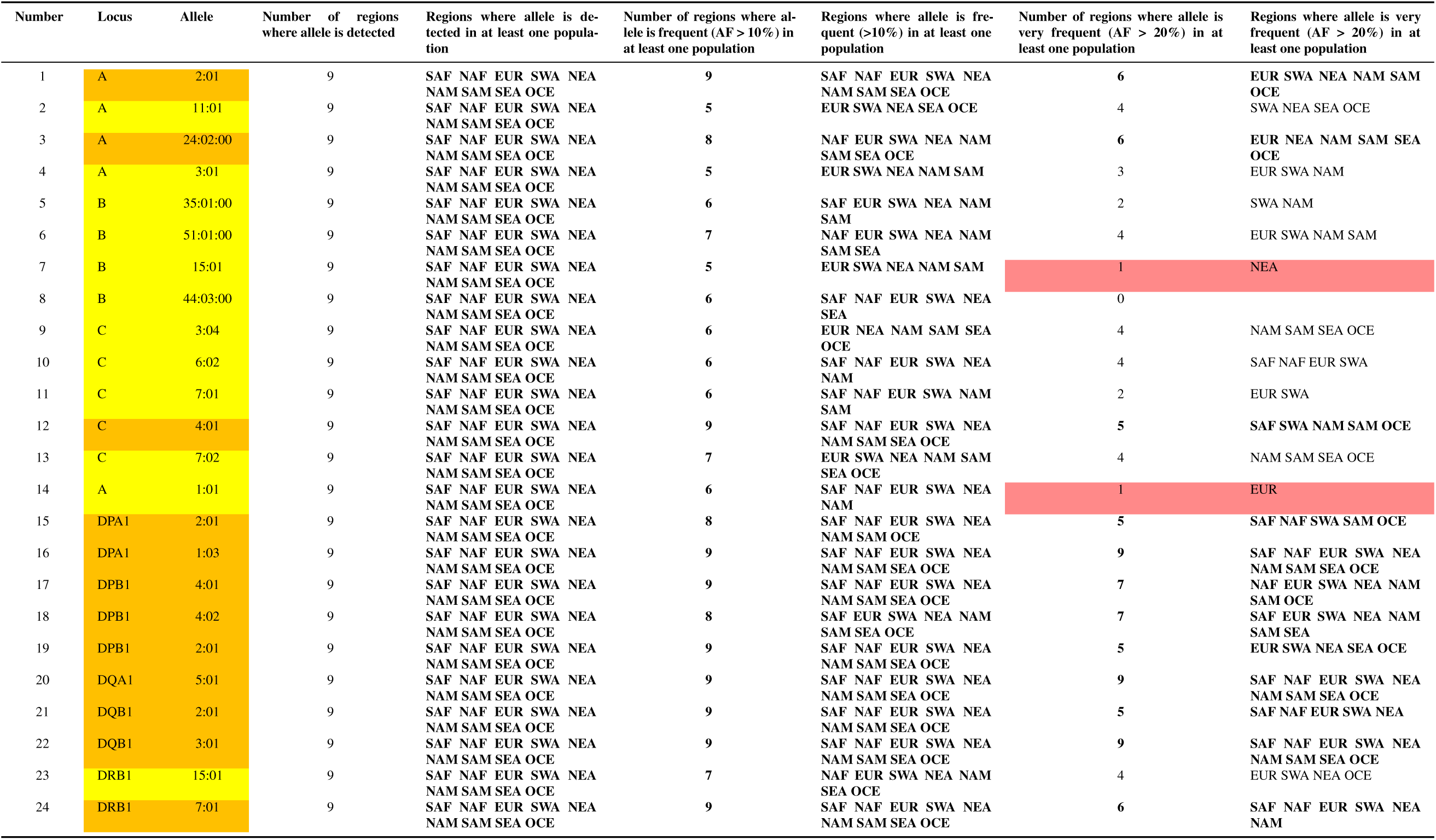
List of HLA alleles with high frequencies in Eurasia. Frequent (allele frequency AF > 10%) or very frequent (allele frequency AF > 20%) alleles (note that very frequent (AF > 20%) alleles are also counted as frequent (AF > 10%) alleles). Regions: SAF: Sub-Saharan Africa; NAF: North Africa; EUR: Europe; SWA: South-West, Central and South Asia; NEA: North-East Asia; NAM: North America; SAM: South America; SEA: South-East Asia; OCE: Oceania (Pacific, New-Guinea & Australia). Yellow: Universally frequent alleles (AF > 10% in at least one population of most regions, i.e. more than 4 regions); Orange: Universally very frequent alleles (AF > 20% in at least one population of most regions, i.e. more than 4 regions); Red: Locally frequent or very frequent (AF > 10% or AF > 20% in a single region). Bold indicates all cases where most regions (i.e. more than 4 regions) are involved. AF: Allele frequency.

**Table S3:**
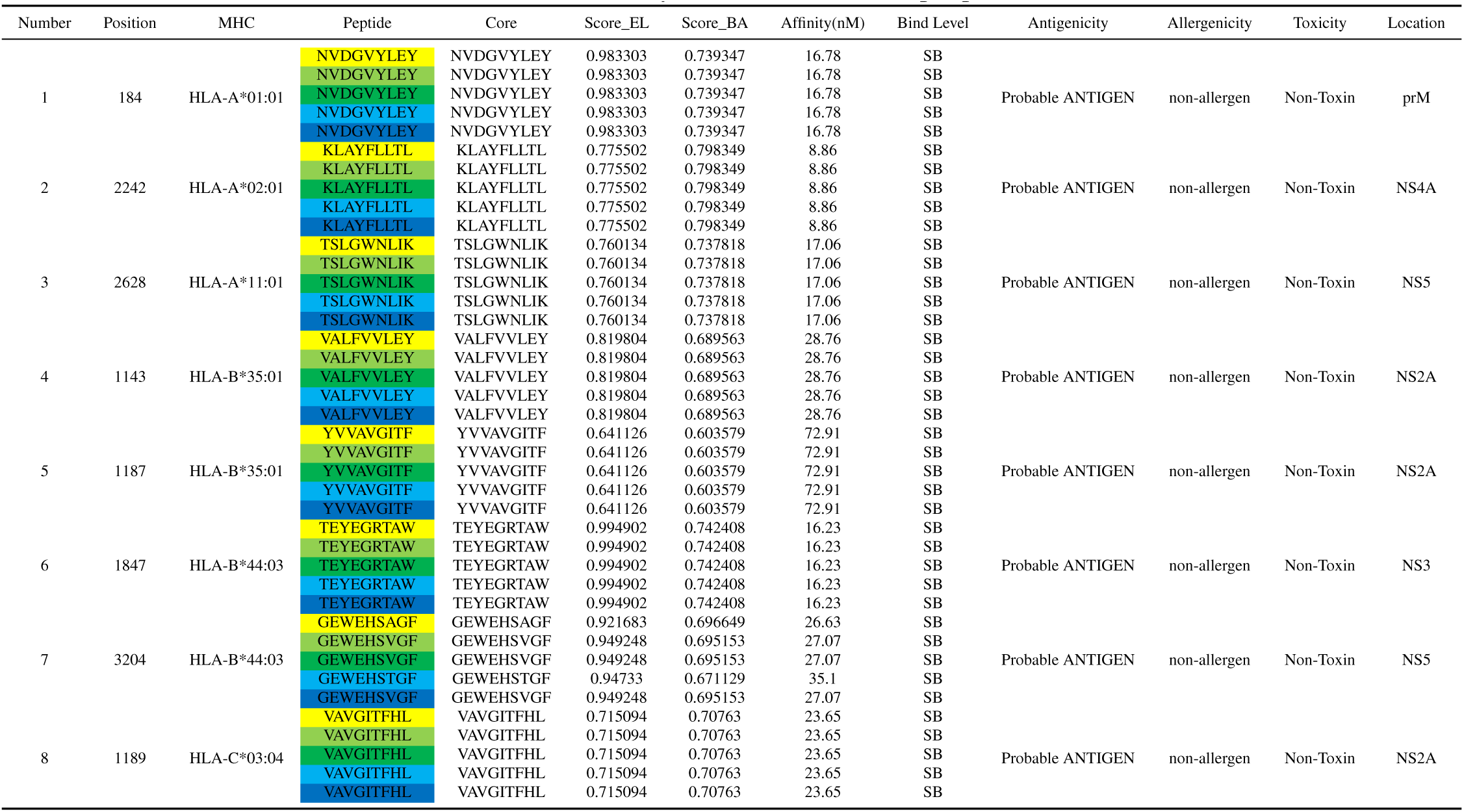
Summary of the screened CTL epitopes.

**Table S4:**
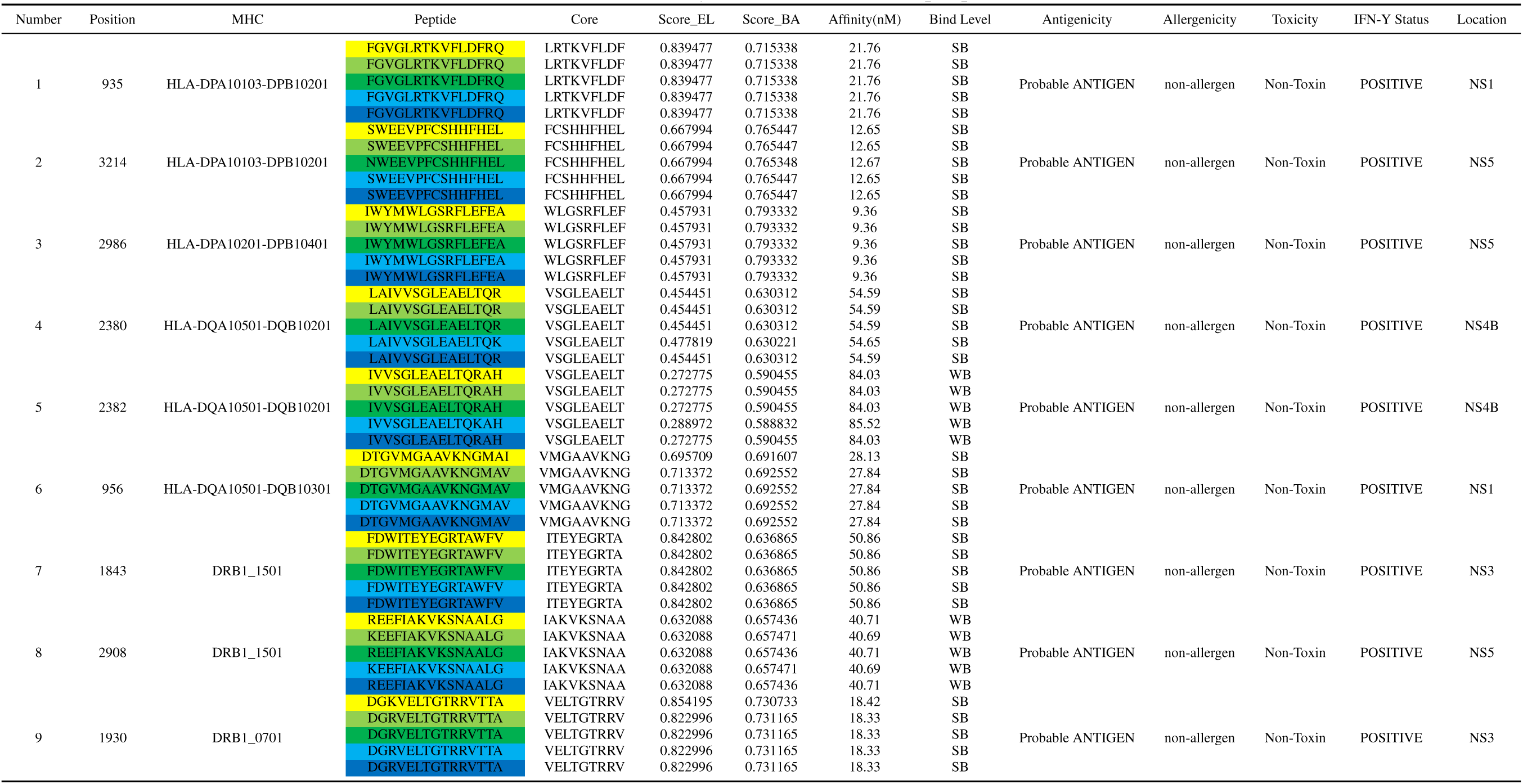
Summary of the screened HTL epitopes

**Table S5:** The population coverage of the selected epitopes across different countries and ethnicities.

| population/area | Class I |  |  | Class II |  |  | Class combined |  |  |
| --- | --- | --- | --- | --- | --- | --- | --- | --- | --- |
|  | coverage | average_hit | pc90 | coverage | average_hit | pc90 | coverage | average_hit | pc90 |
| Belgium | 73.70% | 0.91 | 0.38 | 90.28% | 1.5 | 1.01 | 97.44% | 2.41 | 1.46 |
| Canada | 0.00% | 0 | 0 | 93.14% | 2.24 | 1.16 | 93.14% | 2.24 | 1.16 |
| Central Africa | 39.10% | 0.46 | 0.16 | 96.11% | 2.42 | 1.28 | 97.63% | 2.88 | 1.52 |
| Central America | 0.00% | 0 | 0 | 85.52% | 1.97 | 0.69 | 85.52% | 1.97 | 0.69 |
| China | 67.14% | 0.87 | 0.3 | 95.28% | 2.41 | 1.24 | 98.45% | 3.28 | 1.88 |
| Czech Republic | 77.24% | 1.08 | 0.44 | 96.51% | 2.62 | 1.36 | 99.21% | 3.71 | 2.21 |
| East Africa | 43.76% | 0.52 | 0.18 | 95.27% | 2.5 | 1.25 | 97.34% | 3.02 | 1.55 |
| East Asia | 62.98% | 0.87 | 0.27 | 77.87% | 1.37 | 0.45 | 91.81% | 2.23 | 1.08 |
| England | 85.54% | 1.25 | 0.69 | 95.25% | 2.3 | 1.22 | 99.31% | 3.55 | 2.19 |
| Europe | 79.24% | 1.13 | 0.48 | 99.94% | 3.31 | 2.15 | 99.99% | 4.44 | 3.02 |
| Finland | 84.11% | 1.3 | 0.63 | 85.55% | 1.24 | 0.69 | 97.70% | 2.54 | 1.51 |
| France | 72.81% | 0.98 | 0.37 | 99.99% | 3.66 | 2.35 | 100.00% | 4.64 | 3.13 |
| Germany | 82.47% | 1.23 | 0.57 | 95.46% | 2.49 | 1.27 | 99.20% | 3.72 | 2.21 |
| Japan | 64.10% | 0.89 | 0.28 | 92.20% | 1.97 | 1.07 | 97.20% | 2.86 | 1.51 |
| North Africa | 52.88% | 0.65 | 0.21 | 91.62% | 2.12 | 1.06 | 96.05% | 2.77 | 1.38 |
| North America | 73.50% | 1.04 | 0.38 | 99.99% | 3.47 | 2.14 | 100.00% | 4.51 | 2.98 |
| Northeast Asia | 67.35% | 0.87 | 0.31 | 95.28% | 2.41 | 1.24 | 98.46% | 3.28 | 1.88 |
| Oceania | 60.14% | 0.74 | 0.25 | 97.06% | 2.82 | 1.48 | 98.83% | 3.57 | 2.09 |
| Russia | 66.73% | 0.92 | 0.3 | 99.91% | 3.64 | 2.36 | 99.97% | 4.56 | 3.07 |
| South Africa | 39.80% | 0.46 | 0.17 | 0.00% | 0 | 0 | 39.80% | 0.46 | 0.17 |
| South America | 46.54% | 0.56 | 0.19 | 97.54% | 2.81 | 1.46 | 98.69% | 3.37 | 1.9 |
| South Asia | 49.41% | 0.58 | 0.2 | 99.69% | 3.21 | 2.11 | 99.84% | 3.79 | 2.4 |
| Southeast Asia | 64.81% | 0.85 | 0.28 | 71.54% | 1.27 | 0.35 | 89.98% | 2.12 | 1 |
| Sweden | 64.18% | 0.78 | 0.28 | 99.97% | 3.26 | 2.16 | 99.99% | 4.04 | 2.64 |
| Switzerland | 3.76% | 0.04 | 0.1 | 0.00% | 0 | 0 | 3.76% | 0.04 | 0.1 |
| United States | 73.63% | 1.05 | 0.38 | 100.00% | 3.7 | 2.35 | 100.00% | 4.74 | 3.19 |
| West Africa | 49.60% | 0.62 | 0.2 | 98.82% | 3.18 | 1.87 | 99.40% | 3.8 | 2.28 |
| West Indies | 59.85% | 0.75 | 0.25 | 56.20% | 1.02 | 0.23 | 82.42% | 1.78 | 0.57 |
| World | 72.33% | 0.99 | 0.36 | 97.53% | 2.77 | 1.47 | 99.32% | 3.76 | 2.23 |

**Table S6:** The evaluation results after vaccine modeling and refinement.

| Model | Z-Score | Overall quality factor | Ramachandran Plot |  |
| --- | --- | --- | --- | --- |
|  |  |  | Residues in most favoured regions | Residues in disallowed regions |
| 1.1 | -7.28 | 85.013 | 88.50% | 1.80% |
| 1.2 | -7.22 | 80.729 | 87.90% | 1.50% |
| 1.3 | -7.4 | 82.493 | 88.80% | 1.80% |
| 1.4 | -7.25 | 76.762 | 87.90% | 2.10% |
| 1.5 | -7.28 | 84.675 | 87.90% | 1.80% |
| 2.1 | -7.92 | 92.896 | 86.40% | 3.90% |
| 2.2 | -7.84 | 85.205 | 85.50% | 3.30% |
| 2.3 | -8.01 | 87.903 | 87.00% | 3.30% |
| 2.4 | -7.82 | 88.767 | 86.40% | 3.30% |
| 2.5 | -7.86 | 88.889 | 85.20% | 3.60% |
| 3.1 | -6.71 | 88.189 | 85.50% | 2.70% |
| 3.2 | -6.85 | 81.675 | 87.30% | 2.70% |
| 3.3 | -6.78 | 82.984 | 87.00% | 3.00% |
| 3.4 | -6.78 | 86.807 | 86.40% | 2.40% |
| 3.5 | -6.74 | 83.947 | 87.00% | 3.00% |
| 4.1 | -6.81 | 91.545 | 88.80% | 0.90% |
| 4.2 | -6.81 | 87.826 | 89.40% | 0.90% |
| 4.3 | -6.83 | 84.874 | 88.20% | 0.90% |
| 4.4 | -6.73 | 86.835 | 89.10% | 0.90% |
| 4.5 | -7.01 | 86.416 | 88.80% | 0.90% |
| 5.1 | -7.07 | 89.737 | 88.50% | 1.50% |
| 5.2 | -7.15 | 89.501 | 88.20% | 1.20% |
| 5.3 | -6.97 | 87.435 | 86.70% | 1.20% |
| 5.4 | -7.06 | 88.482 | 87.90% | 1.50% |
| 5.5 | -7.07 | 89.791 | 88.50% | 1.20% |

**Table S7:**
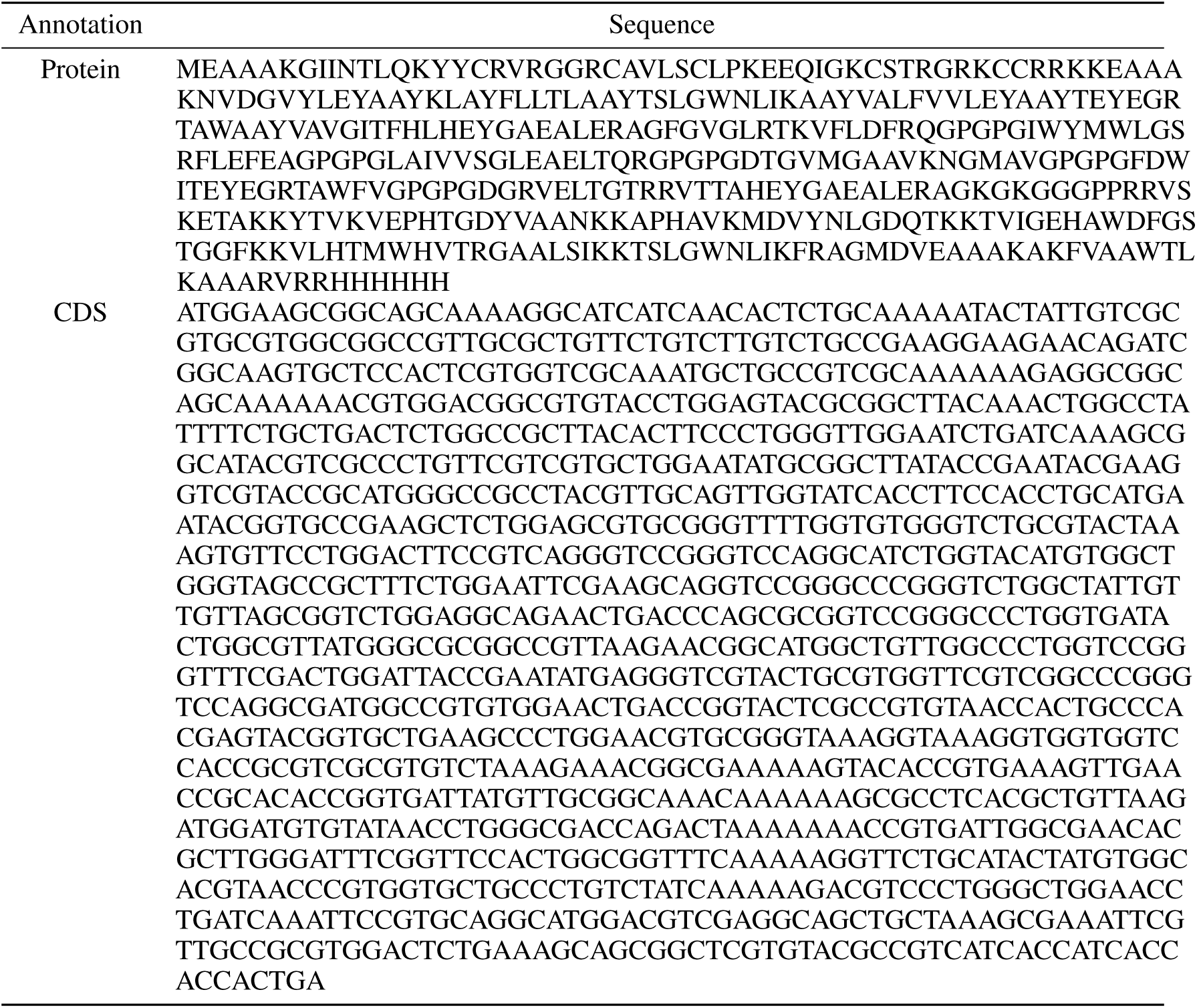
The sequence information of the multi-epitope vaccine.

## References

1. Agudelo, M., Palus, M., Keeffe, J. R., Bianchini, F., Svoboda, P., Salát, J., Peace, A., Gazumyan, A., Cipolla, M., Kapoor, T., et al. (2021). Broad and potent neutralizing human antibodies to tick-borne flaviviruses protect mice from disease. Journal of Experimental Medicine, 218(5):e20210236.

2. Alharbi, M., Alshammari, A., Alsabhan, J. F., Alzarea, S. I., Alshammari, T., Alasmari, F., and Alasmari, A. F. (2024). A novel vaccine construct against zika virus fever: insights from epitope-based vaccine discovery through molecular modeling and immunoinformatics approaches. Frontiers in Immunology, 15:1426496.

3. Andersson, C. R., Vene, S., Insulander, M., Lindquist, L., Lundkvist, Å., and Günther, G. (2010). Vaccine failures after active immunisation against tick-borne encephalitis. Vaccine, 28(16):2827–2831.

4. Arai, R., Ueda, H., Kitayama, A., Kamiya, N., and Nagamune, T. (2001). Design of the linkers which effectively separate domains of a bifunctional fusion protein. Protein engineering, 14(8):529–532.

5. Avšič-Županc, T., Poljak, M., Matičič, M., Radšel-Medvešček, A., LeDuc, J., Stiasny, K., Kunz, C., and Heinz, F. (1995). Laboratory acquired tick-borne meningoencephalitis: characterisation of virus strains. Clinical and diagnostic Virology, 4(1):51–59.

6. Chiffi, G., Grandgirard, D., Leib, S. L., Chrdle, A., and Růžek, D. (2023). Tick-borne encephalitis: A comprehensive review of the epidemiology, virology, and clinical picture. Reviews in medical virology, 33(5):e2470.

7. Dai, X., Shang, G., Lu, S., Yang, J., and Xu, J. (2018). A new subtype of eastern tick-borne encephalitis virus discovered in qinghai-tibet plateau, china. Emerging microbes & infections, 7(1):1–9.

8. De Graaf, J. A., Reimerink, J. H., Voorn, G. P., bij de Vaate, E. A., De Vries, A., Rockx, B., Schuitemaker, A., and Hira, V. (2016). First human case of tick-borne encephalitis virus infection acquired in the netherlands, july 2016. Eurosurveillance, 21(33):30318.

9. Dhanda, S. K., Vir, P., and Raghava, G. P. (2013). Designing of interferon-gamma inducing mhc class-ii binders. Biology direct, 8:1–15.

10. Dimitrov, I., Flower, D. R., and Doytchinova, I. (2013). Allertop-a server for in silico prediction of allergens. In BMC bioinformatics, volume 14, pages 1–9. Springer.

11. Doytchinova, I. A. and Flower, D. R. (2007). Vaxijen: a server for prediction of protective antigens, tumour antigens and subunit vaccines. BMC bioinformatics, 8:1–7.

12. Füzik, T., Formanová, P., Růžek, D., Yoshii, K., Niedrig, M., and Plevka, P. (2018). Structure of tick-borne encephalitis virus and its neutralization by a monoclonal antibody. Nature communications, 9(1):436.

13. Gupta, S., Kapoor, P., Chaudhary, K., Gautam, A., Kumar, R., Consortium, O. S. D. D., and Raghava, G. P. (2013). In silico approach for predicting toxicity of peptides and proteins. PloS one, 8(9):e73957.

14. Hebditch, M., Carballo-Amador, M. A., Charonis, S., Curtis, R., and Warwicker, J. (2017). Protein–sol: a web tool for predicting protein solubility from sequence. Bioinformatics, 33(19):3098–3100.

15. Heo, L., Park, H., and Seok, C. (2013). Galaxyrefine: Protein structure refinement driven by side-chain repacking. Nucleic acids research, 41(W1):W384–W388.

16. Ikeda, M., Arai, M., Lao, D. M., and Shimizu, T. (2002). Transmembrane topology prediction methods: a reassessment and improvement by a consensus method using a dataset of experimentally-characterized transmembrane topologies. In silico biology, 2(1):19–33.

17. Im, J. H., Baek, J.-H., Durey, A., Kwon, H. Y., Chung, M.-H., and Lee, J.-S. (2020). Geographic distribution of tick-borne encephalitis virus complex. Journal of Vector Borne Diseases, 57(1):14–22.

18. Jiménez-García, B., Pons, C., and Fernández-Recio, J. (2013). pydockweb: a web server for rigid-body protein–protein docking using electrostatics and desolvation scoring. Bioinformatics, 29(13):1698–1699.

19. Karypis, G. (2006). Yasspp: better kernels and coding schemes lead to improvements in protein secondary structure prediction. *Proteins: Structure*, Function, and Bioinformatics, 64(3):575–586.

20. Kim, D. E., Chivian, D., and Baker, D. (2004). Protein structure prediction and analysis using the robetta server. Nucleic acids research, 32(suppl_2):W526–W531.

21. Kollaritsch, H., Paulke-Korinek, M., Holzmann, H., Hombach, J., Bjorvatn, B., and Barrett, A. (2012). Vaccines and vaccination against tick-borne encephalitis. Expert review of vaccines, 11(9):1103–1119.

22. Kwasnik, M., Rola, J., and Rozek, W. (2023). Tick-borne encephalitisreview of the current status. Journal of Clinical Medicine, 12(20):6603.

23. Laskowski, R., MacArthur, M., and Thornton, J. (2006). Procheck: validation of protein-structure coordinates.

24. Ličková, M., Fumačová Havlíková, S., Sláviková, M., and Klempa, B. (2021). Alimentary infections by tick-borne encephalitis virus. Viruses, 14(1):56.

25. Lindquist, L. and Vapalahti, O. (2008). Tick-borne encephalitis. The Lancet, 371(9627):1861–1871.

26. Nielsen, H. (2017). Predicting secretory proteins with signalp. Protein function prediction: methods and protocols, pages 59–73.

27. Paz, J. O., Batchelor, W. D., and Pedersen, P. (2004). Webgro: A web-based soybean management decision support system. Agronomy Journal, 96(6):1771–1779.

28. Pierson, T. C. and Diamond, M. S. (2020). The continued threat of emerging flaviviruses. Nature microbiology, 5(6):796–812.

29. Pulkkinen, L. I., Butcher, S. J., and Anastasina, M. (2018). Tick-borne encephalitis virus: a structural view. Viruses, 10(7):350.

30. Rani, N. A., Robin, T. B., Prome, A. A., Ahmed, N., Moin, A. T., Patil, R. B., Sikder, M. N. A., Bappy, M. N. I., Afrin, D., Hossain, F. M. A., et al. (2024). Development of multi epitope subunit vaccines against emerging carp viruses cyprinid herpesvirus 1 and 3 using immunoinformatics approach. Scientific Reports, 14(1):11783.

31. Rapin, N., Lund, O., Bernaschi, M., and Castiglione, F. (2010). Computational immunology meets bioinformatics: the use of prediction tools for molecular binding in the simulation of the immune system. PloS one, 5(4):e9862.

32. Reynisson, B., Alvarez, B., Paul, S., Peters, B., and Nielsen, M. (2020). Netmhcpan-4.1 and netmhciipan-4.0: improved predictions of mhc antigen presentation by concurrent motif deconvolution and integration of ms mhc eluted ligand data. Nucleic acids research, 48(W1):W449–W454.

33. Saha, S. and Raghava, G. P. S. (2006). Prediction of continuous b-cell epitopes in an antigen using recurrent neural network. *Proteins: Structure*, Function, and Bioinformatics, 65(1):40–48.

34. Sanchez-Mazas, A., Nunes, J. M., Dominguez, E. A., Gerbault, P., Faye, N. K., Almawi, W., Andreani, M., Arrieta-Bolanos, E., Augusto, D. G., Buhler, S., et al. (2024). The most frequent hla alleles around the world: A fundamental synopsis. Best Practice & Research Clinical Haematology, page 101559.

35. Sarkar, B., Ullah, M. A., Araf, Y., Das, S., and Hosen, M. J. (2021). Blueprint of epitope-based multivalent and multipathogenic vaccines: targeted against the dengue and zika viruses. Journal of Biomolecular Structure and Dynamics, 39(18):6882–6902.

36. Sayers, E. W., Bolton, E. E., Brister, J. R., Canese, K., Chan, J., Comeau, D. C., Connor, R., Funk, K., Kelly, C., Kim, S., et al. (2022). Database resources of the national center for biotechnology information. Nucleic acids research, 50(D1):D20–D26.

37. Simmonds, P., Becher, P., Bukh, J., Gould, E. A., Meyers, G., Monath, T., Muerhoff, S., Pletnev, A., Rico-Hesse, R., Smith, D. B., et al. (2017). Ictv virus taxonomy profile: Flaviviridae. Journal of General Virology, 98(1):2–3.

38. Sukhorukov, G. A., Paramonov, A. I., Lisak, O. V., Kozlova, I. V., Bazykin, G. A., Neverov, A. D., and Karan, L. S. (2023). The baikal subtype of tick-borne encephalitis virus is evident of recombination between siberian and far-eastern subtypes. PLoS Neglected Tropical Diseases, 17(3):e0011141.

39. Süss, J. (2011). Tick-borne encephalitis 2010: Epidemiology, risk areas, and virus strains in europe and asiaan overview. Ticks and tick-borne diseases, 2(1):2–15.

40. Tarrahimofrad, H., Rahimnahal, S., Zamani, J., Jahangirian, E., and Aminzadeh, S. (2021). Designing a multi-epitope vaccine to provoke the robust immune response against influenza a h7n9. Scientific Reports, 11(1):24485.

41. Thumuluri, V., Martiny, H.-M., Almagro Armenteros, J. J., Salomon, J., Nielsen, H., and Johansen, A. R. (2022). Netsolp: predicting protein solubility in escherichia coli using language models. Bioinformatics, 38(4):941–946.

42. Tonteri, E., Kipar, A., Voutilainen, L., Vene, S., Vaheri, A., Vapalahti, O., and Lundkvist, Å. (2013). The three subtypes of tick-borne encephalitis virus induce encephalitis in a natural host, the bank vole (myodes glareolus). PloS one, 8(12):e81214.

43. Van Zundert, G., Rodrigues, J., Trellet, M., Schmitz, C., Kastritis, P., Karaca, E., Melquiond, A., van Dijk, M., De Vries, S., and Bonvin, A. (2016). The haddock2. 2 web server: user-friendly integrative modeling of biomolecular complexes. Journal of molecular biology, 428(4):720–725.

44. Vita, R., Mahajan, S., Overton, J. A., Dhanda, S. K., Martini, S., Cantrell, J. R., Wheeler, D. K., Sette, A., and Peters, B. (2019). The immune epitope database (iedb): 2018 update. Nucleic acids research, 47(D1):D339–D343.

45. Weng, G., Wang, E., Wang, Z., Liu, H., Zhu, F., Li, D., and Hou, T. (2019). Hawkdock: a web server to predict and analyze the protein–protein complex based on computational docking and mm/gbsa. Nucleic acids research, 47(W1):W322–W330.

46. Wiederstein, M. and Sippl, M. J. (2007). Prosa-web: interactive web service for the recognition of errors in three-dimensional structures of proteins. Nucleic acids research, 35(suppl_2):W407–W410.

